# Poor prognosis in IBD-complicated colon cancer through gut dysbiosis-related immune response failure

**DOI:** 10.1101/2025.03.25.645177

**Authors:** Iradj Sobhani, Nilmara De Oliveira Alves, Mohammad Sadeghi, Cecile Charpy, Emma Bergsten, Aurélien Amiot, Caroline Barau, Francesco Brunetti, Amaury Vaysse, Christophe Tournigand, Mathias Chamaillard, Khashayarsha Khazaie, Denis Mestivier

**Affiliations:** Service de Gastroenterologie, APHP, Hopital Henri Mondor, F-94010 Créteil-France; EA7375 – EC2M3 Université Paris Est Créteil (UPEC); F-94000 Créteil France; ONCOLille, Universite de Lille, Inserm, U1003, Lille, France; Service d’Anatomie pathologique, APHP, Hopital Henri Mondor, F-Créteil-94010 France; Plateforme de Ressources Biologiques, APHP, Hopital Henri Mondor, F-94010 Créteil-France; Service de Chirurgie digestive, APHP, Hopital Henri Mondor, F-94010 Créteil-France; Institut Pasteur, Université Paris Cité, Bioinformatics and Biostatistics Hub, Paris-France; Service d’oncologie médicale, AP-HP, Hopital Henri Mondor, F-94010 Creteil, France; Department of Immunology & Cancer Biology, Mayo Clinic, AZ USA

**Keywords:** Colorectal-Cancer, cardinal-tumor-associated-genes, PHLPP1, microbiota, Immune cells

## Abstract

**Background:** Colorectal cancer (CRC) results from the accumulation of mutations and epigenetic changes in gut epithelial cells likely due to gut microbiota dysbiosis. However, limited research has been done to explore the link between host tumour dysbiosis and disease outcome.

**Methods:** The mechanisms influencing outcomes of 97 colorectal cancer (CRC) patients, including 13 with Lynch syndrome, 20 with inflammatory bowel disease (IBD), and 64 sporadic cases, were analyzed using a multiomics approach. These patients were categorized into two groups: “disease-free/stable disease” and “progression disease” survival outcomes. The analysis included tumor adherent microbiota composition (16S rRNA), somatic gene mutations (WES), gene expression (RNAseq), immune markers (RNAscope), and immune infiltrate cells (immunohistochemistry).

**Results:** IBD-CRC patients had worse outcomes than those with Lynch or sporadic CRC, regardless of TNM staging or treatment. Symbiotic bacteria like *Lactococcus lactis* were significantly reduced in IBD-CRC tissues. Patient outcomes were influenced by the abundance of virulent (*Escherichia coli*) relative to beneficial bacteria (*Lactococcus lactis*). Although no significant increase in deleterious somatic mutations was found in IBD-CRC. 16sRNA revealed increased virulent- and decreased anti-inflammatory symbiotic-bacteria correlating with the upregulation of oncogenes and downregulation of anti-oncogenes like PHLPP1. The multiplex *in situ* hybridization of CD8, IFNγ and PHLPP1 an anti-oncogene revealed significant decrease of immune cells with detectable PHLPP1 expression in IBD-CRC tumour tissues as compared to sporadic CRCs.

**Conclusion:** The poor outcomes in IBD-CRC patients are likely due to gut dysbiosis and immune cell alterations, possibly triggered by microbiota-related epigenetic pathways.

**What You Need to Know:** *BACKGROUND AND CONTEXT:* Colorectal cancer (CRC) is associated with gut microbiota dysbiosis. Inflammatory bowel disease-related CRC (IBD-CRC) is classified as an environment-related condition.

*NEW FINDINGS:* In relation with patient outcomes, tumour tissues from three types of CRC (Sporadic-, IBD-, and Lynch syndrome-CRC) were analyzed using a multiomic approach. This included examining tissue adherent virulent bacteria, gene analyses, and quantifying immune cell infiltration in the mucosa. IBD-CRC patients had the worst outcomes, associated with the down regulation of PHLPP1 gene, virulent/symbiotic imbalance, and immune response failure.

*LIMITATIONS:* Lack of animal experiments using FMT of fresh stool from IBD-CRC patients.

*CLINICAL AND TRANSLATIONAL RESEARCH RELEVANCE:* Among the different types of CRC, IBD-CRC patients showed a greater imbalance between harmful and beneficial bacteria, along with immune response failure.

*Lay summary:* This study compares the pathological and clinical characteristics of patients with colorectal cancer (CRC) across three distinct etiologies: sporadic CRC, inflammatory bowel disease (IBD)-associated CRC, and Lynch syndrome-associated CRC (LS-CRC). Distinct differences in tumor-adherent microbiota, gene expression and immune response profiles were observed. Notably, IBD-CRC patients demonstrated the poorest prognosis depending on microbe-host gene interaction highlighting potential biomarkers for disease prognosis and treatment strategies.

## Introduction

Colorectal cancer (CRC) is one of the most common cancers with elevated deaths per year^1^. Genetic mutations and epigenetic events contribute to cancer susceptibility, particularly in environment-related CRCs (i.e., sporadic, inflammatory bowel disease (IBD))^2^.

Bacteria play key pathophysiological roles in the etiology of CRC^6^, by activating inflammatory cells and modulating immune responses^,5,6^. In CRC, microbiota imbalances lead to an increase in harmful bacteria (pathobionts) and a decrease in beneficial bacteria (symbionts)^7^.However, it remains unclear how these changes affect different CRC types and influence therapy responses and disease outcomes^8^. Additionally, the relationship between bacteria composition, host gene responses, and CRC outcomes are not yet fully understood. Gut microbiota imbalances are thought to increase the risk of CRC in IBD patients, likely due to ongoing chronic inflammation, even when the disease is clinically managed. Thus, our study aimed to investigate how bacterial mediate host DNA changes and expression of genes in CRC tumours. We utilized 16r sRNA, whole exome (WES) and RNA (RNAseq) sequencing as well as immune cells quantification in a large cohort of CRC cases, including IBD patients with cancer occurring in inflammatory colonic or rectal tissues.

We found that CRC tumors are enriched with virulent bacteria, leading to an imbalance with symbionts. This imbalance is associated with increased recruitment of inflammatory cells, alongside a failure of immune cell function and reduced expression of anti-oncogenes like PHLPP1, especially in cases of IBD-related CRC.

## Patients, Materials and Methods

### Patients

For the present study patients referred to University hospitals for colonoscopy and enrolled in several cohort prospective surveys (Acronyms CCR1, Valihybritest, Vatnimad; for description see Ref 5 and ClinicalTrials.gov: NCT01270360), we analyzed materials obtained routinely in symptomatic (CCR1&Valihybritest studies) or asymptomatic patients (Vatnimad from mass screening programs) presenting with an invasive adenocarcinoma at colonoscopy. All patients were included after informed consent and materials obtained before treatment of the tumour. Studies were financed by French government (INCA, PHRC, ANR announcement) and protocols have been approved by the ethics committee (Comité de Protection des Personnes Ile de France IX, no. 10-006, 15/02/2010); clinical, colonoscopy, pathology files were registered.

Primary tumours were endoscopically (stage I) or surgically (stage II, III and two IV cases) removed depending on tumour size and extension; materials were submitted to pathology and molecular analyses; when a patient didn’t undergo surgery (most stage IV CRC) significant samples from primary tumour or metastases (liver) were obtained and similarly analyzed. Standard chemotherapy protocols associated with radiotherapy in neo adjuvant way was given only in Rectal cancer patients. Only CRC patients with stage III and IV (metastatic dissemination) received chemotherapy and/or immune targeted therapy, according to international consensus protocols. No patient with stage I or II received adjuvant chemotherapy until a recurrence was observed. The evaluation to confirm the remission or the progression of the disease was based on a physical examination and blood tests (blood cell counts, liver function tests, and tumour markers), performed 4–5 months after curative-intent surgery in stage II to stage IV and repeated every 6 months up to a period of 3 yrs follow up or death. During this period, 3-to 6-monthly visit and body CT scan imaging was scheduled until recurrence or progression was observed. A total of 97 CRC patients were recruited for the present investigation.

### Histopathology staining

Tumour and normal neighbor tissues were used for pathology diagnosis after hematoxylin and eosin (H&E) staining, for Immunohistochemistry (IHC) and RNAscope *in situ* hybridization on FFPE 4µm thickness tissue sections by using appropriate antibodies (Sigma Aldrich, Invitrogen, France) and RNAscope multiplex fluorescent reagent 2.5 HD kit assay (Advanced Cell Diagnostics, Newark, CA, USA) according to the manufacturer’s instructions. IHC sections were revealed using DAB technique.

### DNA and RNA Isolation and Quantitative PCR

Tissue samples were homogenized with QIA shredders prior to DNA and RNA isolation using the AllPrep DNA/RNA mini kit (Qiagen). RNA (1 ⍰g) was reverse transcribed using a cDNA Reverse Transcription Kit (Applied Biosystems), cDNAs so generated were triplicated and analyzed using SYBR-Green PCR mix (Applied Biosystems) and values were compared to GAPDH expression for normalization. Following primers were used: GAPDH (Forward 5’-GAGAGACCCTCACTGCTG-3’, Reverse 5’-GATGGTACATGACAAGGTGC-3’), PHLPP1 (Forward 5’-ACTGGGATTTGGGGAGCTG-3’, Reverse 5’-CGTCTTGTCCATCGGTTCACT-3’). Quantitative PCR experiments were performed according to methods described elsewhere^4^; briefly Fast Real-time PCR system (Applied Biosystems) and relative mRNA concentrations were calculated by the 2^-ΔΔCt^ method, where Ct is the mean threshold cycle value and GAPDH was used to normalize. Fold changes in gene expression for each sample were calculated using the 2^−ΔΔCq^ method relative to control after normalization of gene-specific Cq values to GAPDH Cq values.

### Microbiota analysis

The 16S rRNA gene sequencing was performed on DNA from frozen tissue samples (extraction using GNOME DNA Isolation Kit-MP Biomedicals) as previously described^9^: after amplification by PCR (V3 to V4 region of the 16S rRNA gene), samples were submitted to a 250-bp paired-end sequencing protocol on the Illumina MiSeq platform. Raw FASTQ files were demultiplexed, quality-filtered using Trimmomatic (sliding windows of 2 with a quality score of 20), and merged using fastq-join from ea-utils (https://expressionanalysis.github.io/ea-utils). Taxonomic assignations were performed using Qiime2 (no quality filtering; default parameters) with the SILVA-123 database; OTU were constructed using UCLUST (threshold of 97% of similarity), Chimera Slayer for chimera removing, and SILVA 16S rRNA database (version 123) for taxonomical assignation. The intergroup high similarity and intragroup low similarity of microbiota were assessed by β-diversity, PCoA (generated by Qiime using unweighted unifrac metrics).

### WES

Samples (Normal/Tumor) were used for mutational analysis [236 (59 x R1/R2 x N/T) fastq files (2×75pb); 29,862,533.2 +/-13,205,899.8 reads/sample; min = 2,840,763; max = 81,265,734] when only tumour samples were submitted to RNAseq [Paired-end reads (2×75pb); 48,200,618.6 +/-4,280,994.8 reads/sample; min= 40,349,619; max = 61,276,451] analysis. The quality was checked using *fastQC* (v0.11.9) and trimmed using the *trimmomatic* software (v0.39, quality threshold Q=20 with a sliding windows of 5 bases and a minimal length of 50pb. Good quality reads for WES were mapped to the *Homo sapiens* genome assembly (GRCh38.95) using the *bowtie2* software (v2.4.4) with default parameters and marked for duplicates using the *Picard* software (v2.26.2). For variant calling, we used the *Varscan* software (v2.2.3, default parameter values except for the purity that was set to 0.4). Variants (SNP and indels) were filtered based on individual p-value < 0.05 and status (“Somatic” / “LOH”). We then used the *snpEff* software (v5.0) to annotate variants, and an in *house python script* to compile statistics. Good quality reads (95.6 +/-0.6 % surviving reads after trimming) for RNAseq were mapped using the STAR software (v2.7.4a) to the *Homo sapiens* genome assembly (GRCh38.95) and we used RSEM (v1.3.1) to calculate gene expression levels. Four out of 97 samples with less than 5M of mapping reads were excluded from further. Two sets of analyses were performed on RNAs. First, we took in consideration all 13361 genes for which reads were reached the threshold for gene identification and analysis in all CRCs and in each type. Then we focused on 137 cardinal genes as dedicated to colon carcinogenesis from the literature (List provided in suppl file) with an FDR cutoff of 0.1. All 137 genes were considered according to “failure (death- or progression disease) vs Success (remission or stable disease)” through an unsupervised analysis. This consisted in a Filtering by Fold Change 1.5, t test [<0.05], and genes which were significantly associated with the failure [All with p = 0.002 and q = 0.186 and without any co factor eliminated when samples withdrawn from the analysis due to very low reads available]. So, active samples for RNA analysis were 95 with normalization Mean=0, and Var=1. According to the median value (3.5) of PHLPP1 RNA, we divided our series into two (High≥3.5 versus Low<3.5) sub groups. Heatmaps were constructed from RNA levels regarding CRC types.

### CMS classification

Three algorithms were applied to RNA values for CMS (Consensus Molecular Subtyping) evaluation using R packages: CMSCaller^10,11^ and CRC Assigner^12^ using default parameters); for each sample, the consensus CMS (CMS most reported by the three callers), was specified and discrepancies reported (see “itmo-RNASEQ-CMS-ALL4packages_rev-09-03-2021-Validated-11-03-2021.ods” in suppl files).

### Gene networks around bacteria

We considered 13361 genes through our RNAseq dataset after filtering threshold of log2(CPM) > 1 gene expression level. We then computed the Pearson correlations between each gene and the PHLPP1 gene for each CRC type. A *p-value* was computed using 1000 permutations. Correlations between a gene RNA and a bacteria number was reported according to p<0.05 and the absolute value of the Pearson coefficient above 0.3 (List provided in suppl file). To construct a network of cardinal genes around a bacteria, correlations were calculated and illustrated using Cytoscape (v3.8.2) with adjusted p<0.05 for multiple analysis.

### RNAscope *in situ* hybridization

The mRNA expression levels for PHLPP1, CD8 and IFNγ were quantified on FFPE tissue sections from CRC tumour tissues using RNAscope multiplex fluorescent reagent 2.5 HD kit assay (Advanced Cell Diagnostics, Newark, CA, USA). The assay was optimized according to the manufacturer’s instructions. Briefly, 5 µm thickness sections were placed on SuperFrost Plus glass slides, were dehydrated, endogenous peroxidases blocked and then, treated with 200 mL of RNAscope target retrieval reagent (1X, 95°C) during 15 min. RNAscope Protease Plus was used to cover each section and slides were placed in HybEZ™ II oven at 40°C for 15 min. The probes hybridization process was performed at 40°C for Hs-IFNγ (310501, Opal 650), Hs-PHLPP1 (1064481-C2, Opal 570) and Hs-CD8A (560391-C3, Opal 520) probes. Between each amplification and staining step, slides were washed twice in 1X RNAscope wash buffer before incubation with DAPI (ThermoFisher Scientific) and treatment with ProLong Gold antifade mounting solution (ThermoFisher Scientific) before imaging. Image acquisition was performed using the Zeiss Axioscan Z1 at 20× magnification through Zen 3.7 software for quantification. This was achieved after specific regions of interest (ROI) were selected on each tissue sample, including tumor regions (n=69 patients) and adjacent normal tissues (n=22 patients). Co-localization analyses of PHLPP1, CD8, and IFNγ stained cells was conducted in two groups categorized by PHLPP1 expression levels (low versus high) and in two (Sporadic, IBD) CRC types. Multiplexed image analysis utilized QuPath 5.0 open-source software^13^, with cell segmentation performed using the Stardist module^14^. The expression of multiplexed markers by these segmented cells was analyzed using artificial intelligence techniques to determine percentage of positive cells.

### Immunohistochemistry characterization of inflammatory and immune cells

To validate RNAScope results, we performed a quantification study on both, tumour and normal neighboring tissues (n=95 CRC cases; 2 excluded due to missing material). Following specific markers were targeted using appropriate antibodies: specific antibodies targeting CD3, CD4, CD8, CD68, CD163, mastocytes, FoxP3, ROR⍰T cells, Ki67, Granzyme were purchased (from Invitrogen, Biotechne and Sigma Aldrich, France) and used according to the company’s instructions and reactions were revealed by DAB. QuPath (https://qupath.github.io/) program version 2.1 was used under windows for quantification; all comparisons were performed using Qlucore program (see statistical section).

### Statistical analyses

The following criteria were applied for all analyses on tissue slides: p value produced by Mann−Whitney−Wilcoxon test after FDR correction < 0.05, fold change > 1 or < −1.

We subjected study populations to packages in Shaman Webserver; these used the DESeq2 for differential expression analyses when statistical analyses have been corrected according to gender, age, and BMI and adjusted for multiple testing. Differential bacteria were then applied to Qlucore program for analyzing tissue adherent bacteria regarding additional associations between CRC type subgroups (sporadic vs LS vs IBD; PHLLP1 high vs low expression).

For patients’ outcomes analyses, characteristics of study populations were described using number (%) for qualitative variables and mean (± SD) for quantitative variables. All patients underwent a survey program of 3 years or more. They have been considered in “failure” or “success” regarding the course the disease as compared to the baseline. The failure was defined when a combined endpoint including death, recurrence after remove of all (primary as well as metastases) tumour locations, or progression disease (stage IV CRC patients who received only chemotherapy) was reached during the follow up period. The probability of the survival (disease free, or no progression considered with death censured) was calculated according to CRC types with significant difference set to adjusted p<0.05. Death or the progression of the disease over at least three years follow up for each patient ranged under Kaplan Meier construction. Omics Explorer Qlucore 3.2 software for survival statistical analysis with gene and bacteria associations was used. In this program, variance filtering is used to reduce the noise, the “Mean=0, Var=1” setting and to scale the data without any variable left after the filtering. The Principal Component Analysis (PCA) was used to visualize the data set in a three-dimensional space. P-values were adjusted for multiple testing using the Benjamini-Hochberg method^21-23^, and variables with adjusted p-values below 0.05 were considered significant. For main differential molecular markers (taxa, genes and genes), two analyses were performed to compare CRC tumour tissue types with and without Bonneferoni correction. Effect on “failure” or “success” in either below or above RNA median levels of gene RNAs associated were investigated using χ2 test or Fisher’s exact test and the Mann–Whitney test when appropriate.

## RESULTS

### Pathology and clinical description of patients and their outcome

Patient [sporadic- (n=64), IBD- (n=20) and Lynch syndrome-CRC (LS-CRC; n=13)] and disease characteristics (Table 1) were balanced in the various CRC types except for age and long-term immunosuppressive treatments. As expected, the IBD-CRC patients were younger than LS-CRC and sporadic cases, and received immunosuppressive drugs (*i*.*e*. anti TNF⍰, derivatives of azathioprine or methotrexate). Otherwise, there were no significant differences between the three CRC types in terms of CRC TNM staging and tumor therapy (surgery, chemotherapy) protocols.

**Table 1:**
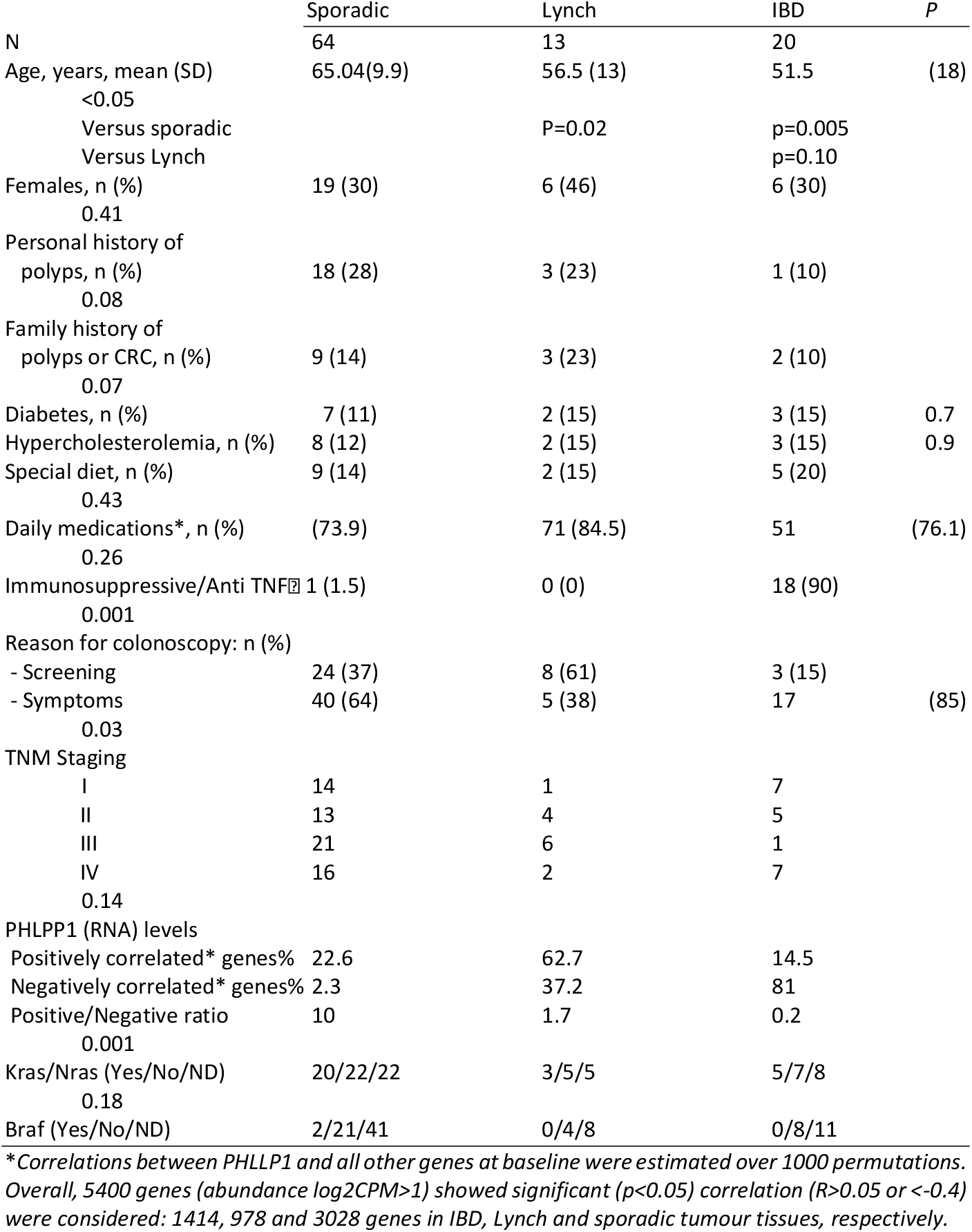
Characteristics of patients according to CRC type.

By using Qlucore model based on filtering out variables with low overall variance to reduce the impact of noise, those patients who experienced “failure” during at least three years follow up were considered according to CRC types; co factors such as age, tumor location (left vs. right), TNM staging (I to IV), that might influence patient’s outcome during the follow-up period were included in the model as co variable. IBD-CRC patients experienced the worse prognosis while LS-CRC patients showed better outcome as compared to those with sporadic CRC or IBD-CRC (Figure 1). We first analyzed tumour tissue adherent bacteria and noticed they differed significantly depending on CRC types.

**Figure 1.**
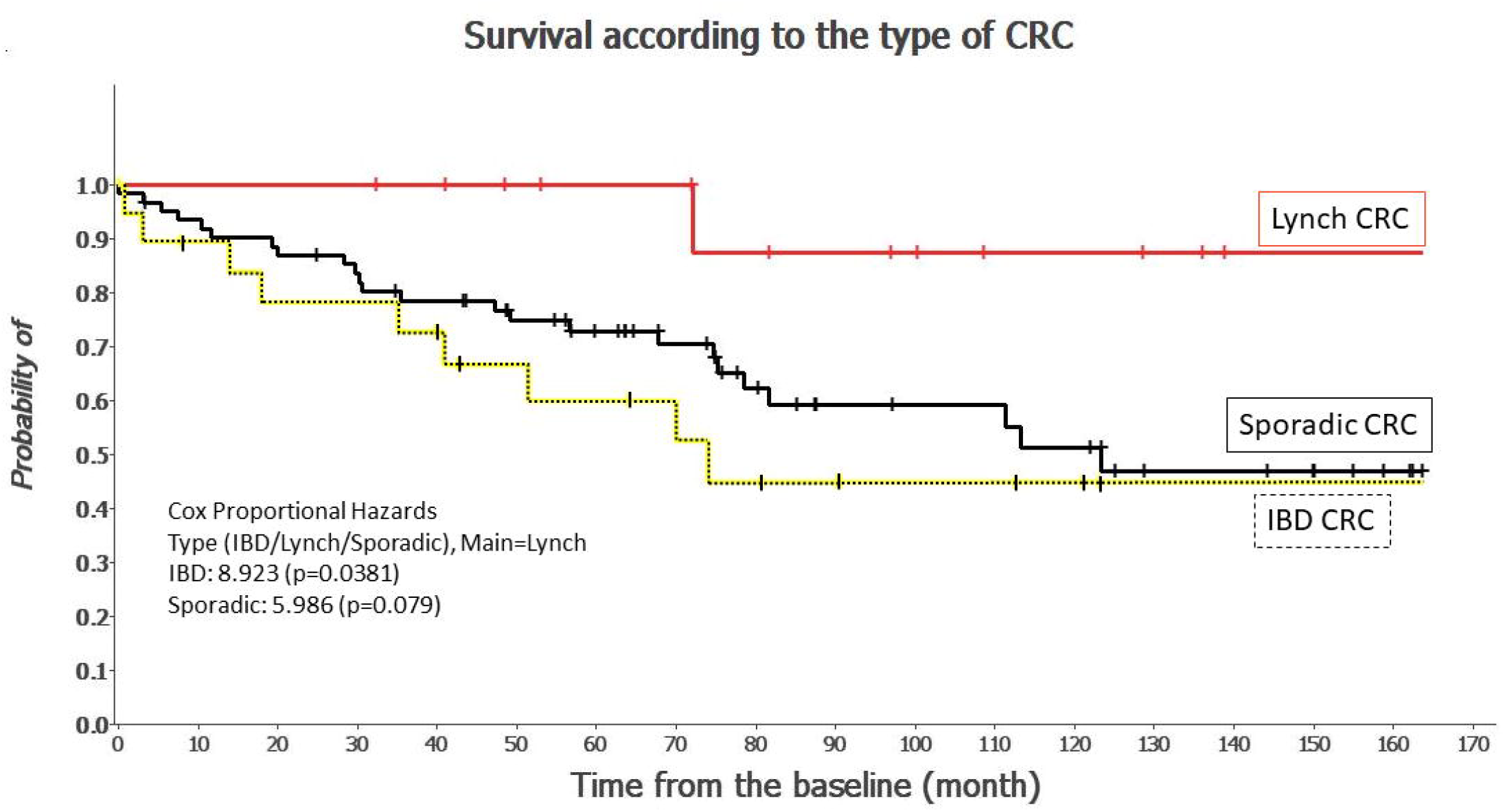
Clinical files of CRC patients including baseline and 3-yr period follow up data, were used to classify each patient in “failure” versus “success”. Dataset was the submitted to Kaplan Meier analysis according to CRC subtypes.

### Tissue adherent microbiota and outcome

PCoA analysis showed significant separations between Sporadic(S)-, Lynch (L)- and IBD(I)-CRC types (Figure 2A). Using Qlucore statistical software analysis and taking all tumour tissue samples together, we observed significant associations between several virulent bacteria adherent to tumour tissues and patients’ outcomes (Figure S1). Again, these bacteria in IBD-CRC patients showed different distribution compared to other tumour types, as illustrated by Krona multilevel pie charts that visualize both the most abundant and their most specific microorganisms (Figure S2). Of note, tumor tissues showed significantly lower *lactococcus lactic* and a trend to the diminution of *Bifidobacterium longum* than normal neighboring tissues (Figure 2B-D; Table S1).

**Figure 2.**
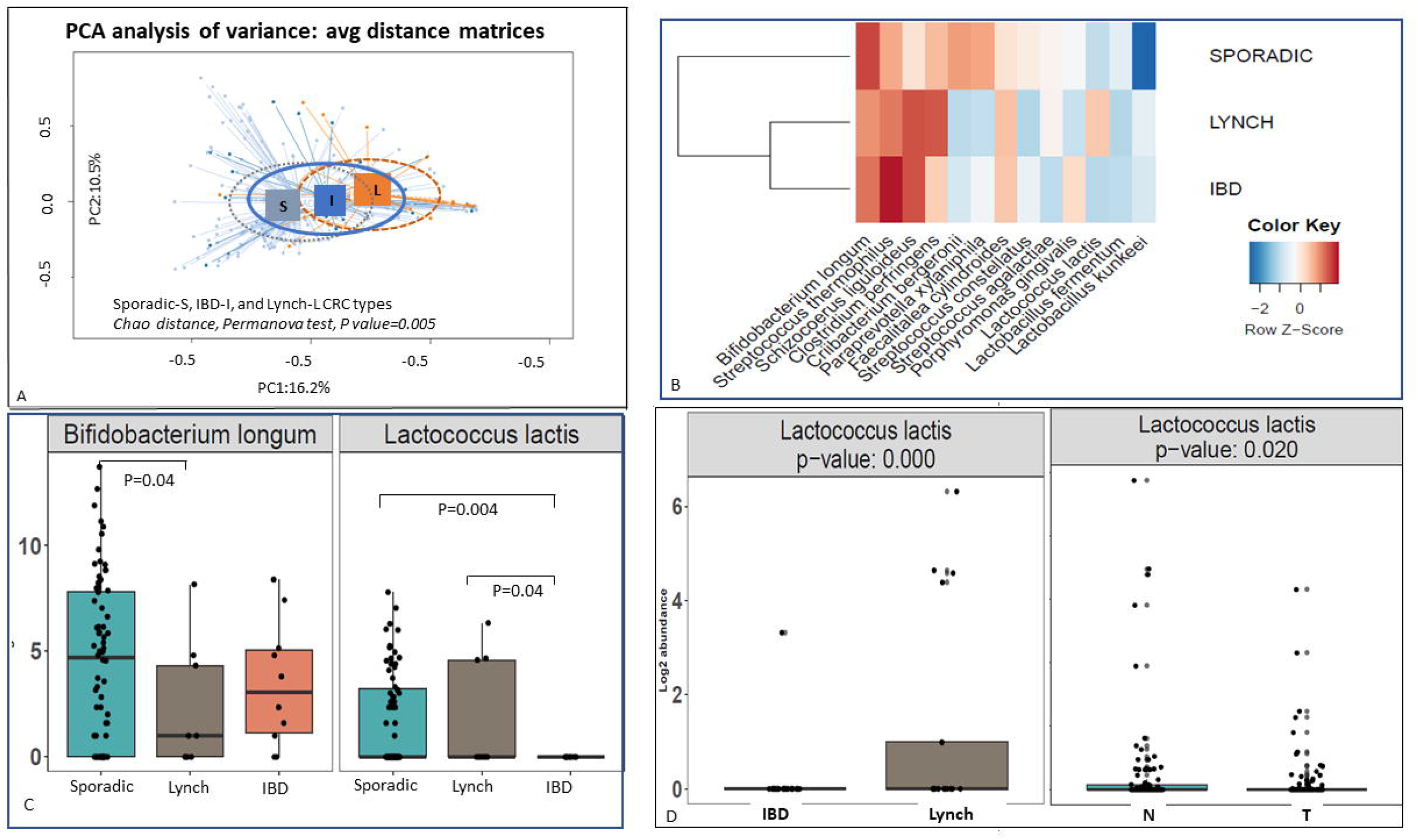
Prokaryote DNA was extracted from all tissue samples and sequences were assigned to bacteria. A. PCoA analysis showed significant separations between Sporadic(S)-, Lynch (L)- and IBD(I)-CRC types. Various virulent and symbiotic genera (B) and various symbiotic species (C) were found significantly different in IBD-CRC tissues. Taking all samples together, tumour (T) adherent Lactococcus lactis was lower as compared to normal neighboring tissues (D).

Higher levels of virulent bacteria such as *porphyromonas gingivalis, shigella bonydii* adherent to tumour tissues were observed in IBD-CRCs contrasting with symbionts (i.e. *Lactococcus lactis*; *Bifidobacterium longum*). For example, *Lactococcus lactis* in IBD tumour tissues was significantly lower than in either Lynch (IBD-CRC/Lynch-CRC 2logFold = −5.394; p=0.04) or sporadic (IBD-CRC/sporadic-CRC 2logFold = −6.574; p=0.004) tumour tissues.

### Pathways associated with co-regulated genes in IBD-CRC

Assuming differential adherent bacteria influence gene up- and down-regulation within tumour tissues differently, we classified each sample in Consensus Molecular Subtypes (CMS) according to Guinney et al., an algorithm modified by Eide et al. ^10,17^ based on RNA values (n=13360). Notably, 60% of IBD-CRC tumours were observed in CMS4 and 35% in CMS3 suggesting EMT and inflammatory/immune cell functional pathways were involved (Table S2, suppl file). Hence, we performed further analyses including both, genes and bacteria. Analyses in our RNA dataset, were based on the statistics of negatively correlated genes in a linear model, and gene RNA levels were ranked through panel of associated genes by using Gene Set Enrichment Analysis (GSEA)^18^. Comprehensive analysis of all 13,360 gene RNAs from tumor tissues across all CRC types, identified 1,576 genes that were significantly associated with at least one other gene.

To enhance the statistical power, we performed a statistical analysis including only 137 cardinal genes (List Table S3 suppl file)) involved in colon carcinogenesis by using Qlucore model based on filtering out variables with low overall variance to reduce the impact of noise. The identification of significantly differential variables (RNA levels) between “failure” and “success”, were considered as predictive of outcome. Thus, PHLPP1 gene together with TRAPP and IL17RB genes were identified as influencing patients’ outcomes with the following covariates: age, tumor location (left vs. right), TNM staging (I to IV), death during the 3 years follow-up period (Figure S3). Further, when these 137 cardinal genes were considered in our series independently from survival, through an inter-gene unsupervised association, several other genes clustered with these three genes (Figure 3).

**Figure 3.**
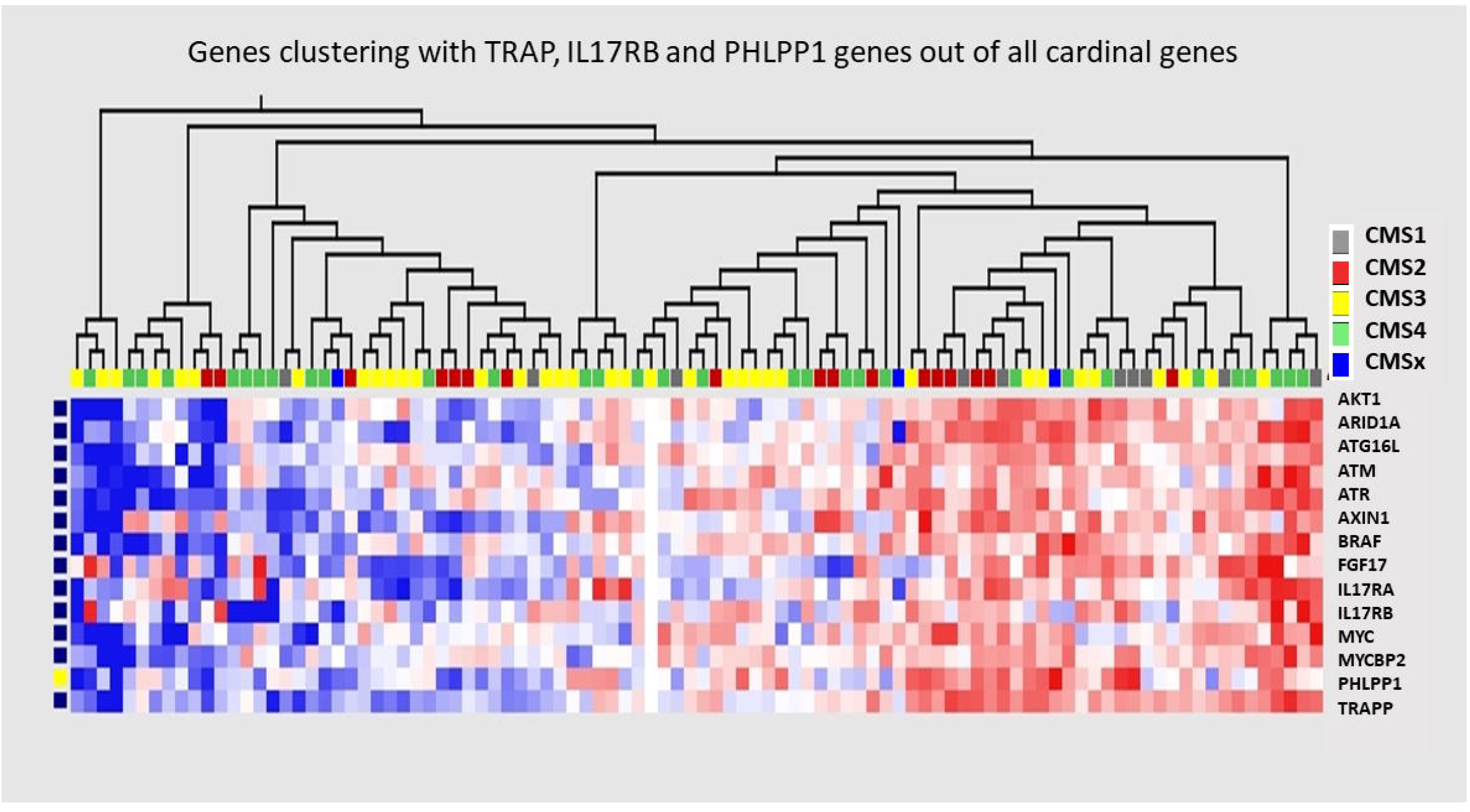
Tumour samples from CRC patients were submitted to RNA extract and gene expression were used for clustering genes through an unsupervised analysis based on 137 cardinal genes; t-test [<] was used comparing gene RNA values according to success (Disease Free or stabile disease over surveillance period) versus Failure (Recurrence/Death or the progression of the disease-PD) using Qlucore Omics Explorer program that enables variance filtering to reduce the noise, and the projection score to set the filtering threshold with 1 vs 0; p = 0.002, q = 0.186 and none eliminated factors filtered by Standard Deviation (s/smax). The heatmap shows levels of gene RNAs regarding CRC types^51^. The identification of significantly differential variables was performed with gene level class as a predictor. P-values were adjusted for multiple testing using the Benjamini-Hochberg method^15^, and variables with adjusted p-values below 0.1 were considered significant.

Amongst, ten genes are involved in autophagy, proliferation, and metabolite traffic, and exhibited significant correlations together in IBD cases. This panel included PHLPP1 gene that was associated with APC (a common initiator of colorectal cancer), FBXW7 (an F-box protein suppressor of JUN and MYC), oncogenes like BRAF, MAP3K3, PDGFRA, FGFR1, PIK3CA and genes related to autophagy and protein traffic (TRAPP), immune-modulating genes (RORC, TLR4, CXCL1, SOCS1), chromatin modifiers (TCF4, TOPRRS), DNA repair genes (PMS2, MLH1, MSH2, MSH6, MGMT), the G2/M cell cycle checkpoint gene MCPH2, and core circadian genes CLOCK and ARMTL2 (BMAL) (Table S4). PHLPP1 and TRAPP/TRAPPC8B genes were significantly down expressed in IBD-CRC tumours as compared to sporadic and LS-CRC tumour tissues (Figure S4A&S4B); all these genes were correlated with AKT1 gene (Figure S4C).

Also, in each CRC subtype (sporadic-, LS- and IBD-CRC), we identified panels of genes whose expression was positively or negatively associated with PHLPP1 gene expression. Six genes (AXIN1, TRRAP, KMT2D, ARID1A, CREBP, SIPA1) positively co-associated with PHLPP1 gene expression in all CRC patients with the most significant correlations having been observed between AKT1, TRAPP and PHLPP1 genes (Figure S5A); remaining co-associations were specific to CRC subtypes with nine genes (BRCA1, CHEK1, CHEK2, MSH6, MSH2, EPCAM, NRAS, DNAJC2, XRCO5) showing a negative co-variation with PHLPP1 gene. Neither significant mutation nor variants in the PHLPP1 gene body were observed in IBD-CRC tissues as assessed by WES analysis of tumour tissue DNAs; while in only 11 out of 96 tissue samples in sporadic (n=9) or Ls (n=2) such nucleotide variations were observed (Table S5).

### Associations between adherent bacteria and host gene expressions

Several adherent species including *E. Coli* and *Shigella bonydii* were found overabundant in tumours than in normal neighboring samples when in tumour tissues *shigella bonydii* and *porhyromaons gingivalis* were differential between sporadic and IBD cases when PHLPP expression (High vs Low) was considered (Figure 4). More generally virulent/symbiotic ratio imbalance in IBD-CRC tumours was noticed as illustrated by *Hemophilus, Porphyromonas, Treponema, Leptotrichia* and *Fusobacterium* genera (Figure S6) and related species (Table S1) which were significantly associated to both tumour location (vs normal neighboring) and CRC types (dramatic imbalance in IBD-CRC cases). Genes positively co-associated with PHLPP1 in CRC-IBD were involved in inflammation (IL17RD, CX3CL1), fibrosis (COL1GA1), and cell survival (FGF1, FGF2, IFG1). Genes negatively co-associated with PHLPP1 in CRC-IBD were cell cycle and DNA repair including BRCA1, CHECK2, and MSH2. LS- and sporadic-CRCs showed positive association of PHLPP1 with cellular oncogenes (CTNNB1/ β-catenin, TP-53, K-ras, ERBB2/HER2, SMAD4), and the autophagy regulatory gene ATG16L1, inflammation (IL17RA), anti-microbial defense (NOD2), replication (ATG), angiogenesis (VEGFA), survival and transformation (EGFR, AKT1, Myc, TOPORS) (Figure S5A). These associations were most significant in IBD-CRC cases. In LS- and sporadic-but not in IBD-CRC cases, we found negative correlations with Methylguanine methyltransferase (MGMT) a DNA repair gene, which is involved in epigenetic regulation of genomic stability (Figure S5A).

**Figure 4.**
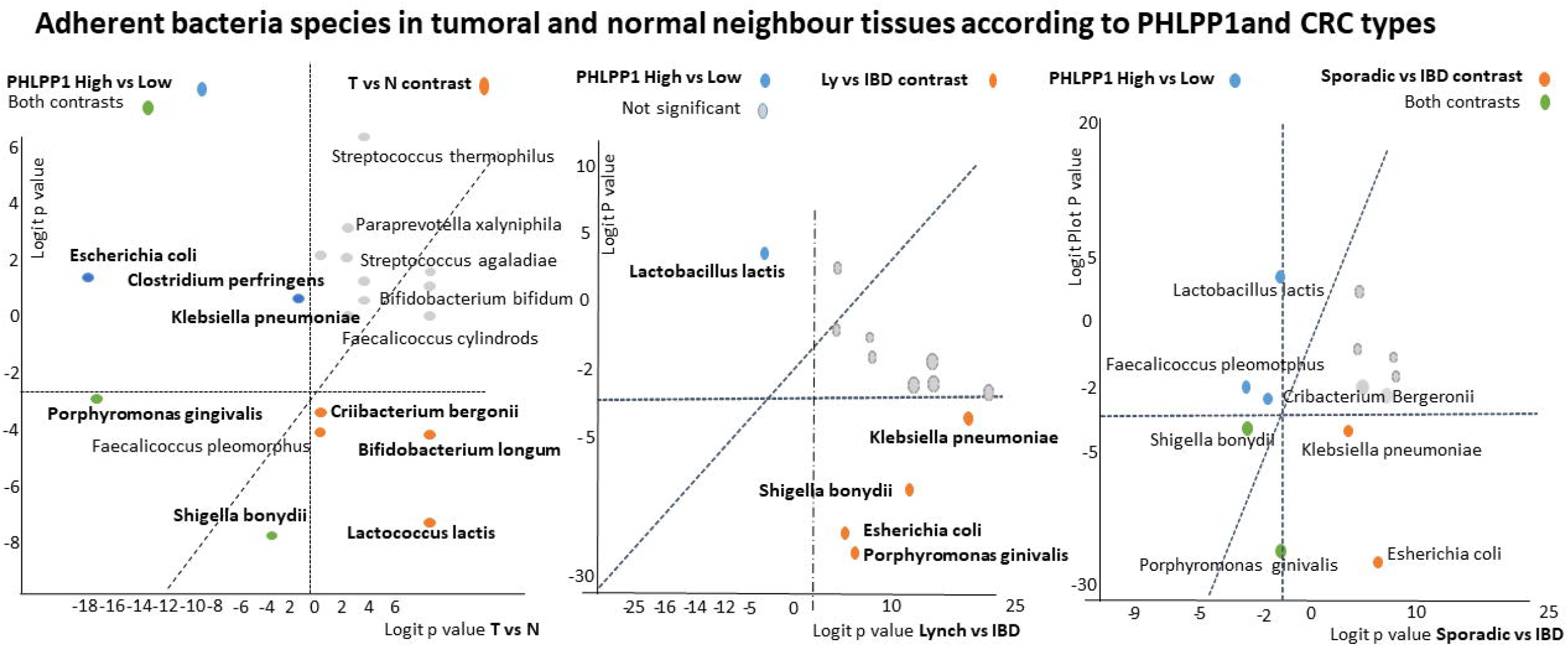
DNA from (T=tumour, N=normal neighbor) tissues were extracted and submitted to 16sRNA sequencing procedure (Illumina technology). Results were analyzed using Shaman platform. Contrasts regarding significant species (contrasts colored by bleu, orange or green) of adherent to tissues are shown depending on CRC types (IBD vs Ly or Sporadic) in orange and on PHLPP1 expression in bleu or both (in green) revealed differential features.

Among 1,576 genes having a significant association with at least another gene (out of all 13,360) across all CRC types in tumor tissues, more than 100 genes correlated with that of PHLPP1 in a manner that was dependent on CRC types. KEGG and EnrichR program^19^ based on cluster rammer analysis was used to gain further insights into the pathways. Infectious Diseases and Cancer Disease were the two main signaling pathways associated with PHLPP1 gene function in the present dataset. Number of genes that were negatively correlated with PHLPP1 were three times higher in IBD-CRC than in sporadic CRC: 30 genes in sporadic-CRCs and 95 genes in IBD-CRCs; no gene was significantly and negatively associated with PHLPP1 in LS tumour tissues. Main pathways identified based on PHLPP1 negatively associated genes in sporadic- and IBD-CRCs, included oxidative phosphorylation, thermogenesis, mitophagy, and proteolysis in sporadic-CRC (Figure S5B). Spliceosome was the major pathway affected in IBD-CRCs highlighting the possible role of PHLPP1 down regulation (Figure S5C) in the splicing complex^20,21^. Hence, from all genome (13360 measurable gene RNAs) that include 137 cardinal genes, we observed.

In order to characterize part of host tissue gene and adherent bacteria interactions in these pathways, and to illustrate virulent/symbiotic associations, we selected bacterium such as *Escherichia* as representative of virulent genera, and *Faecalibacteriul* and *Bifidobacterium* as representative of symbiotics. Then, we showed positive and negatives correlations between these bacteria and genes. We observed more bacteria-gene correlations in IBD than in Sporadic cases whatever all genome (Figure 5B) or only cardinal genes (Figure 5C) were considered; there were more associations between Escherichia and cardinal genes in IBD than in sporadic cases (Figure 5A). While higher numbers of negative correlations were observed between symbiotic bacteria (*Faecalibacterium, Bifidobacterium*), and genes in sporadic-CRC tissues than in IBD-CRCs (Figure 5B). *Faecalibacterium* was found positively associated with RORC, a regulatory gene of immune response in sporadic-but not in IBD-CRC cases (Figure 5C). Balance between *Escherichia Coli* and *Faecalibacterium Praunitzii* species adherent to tissues revealed differential features in IBD- and other (sporadic- or LS-) tumour tissues (Figure 5D).

**Figure 5.**
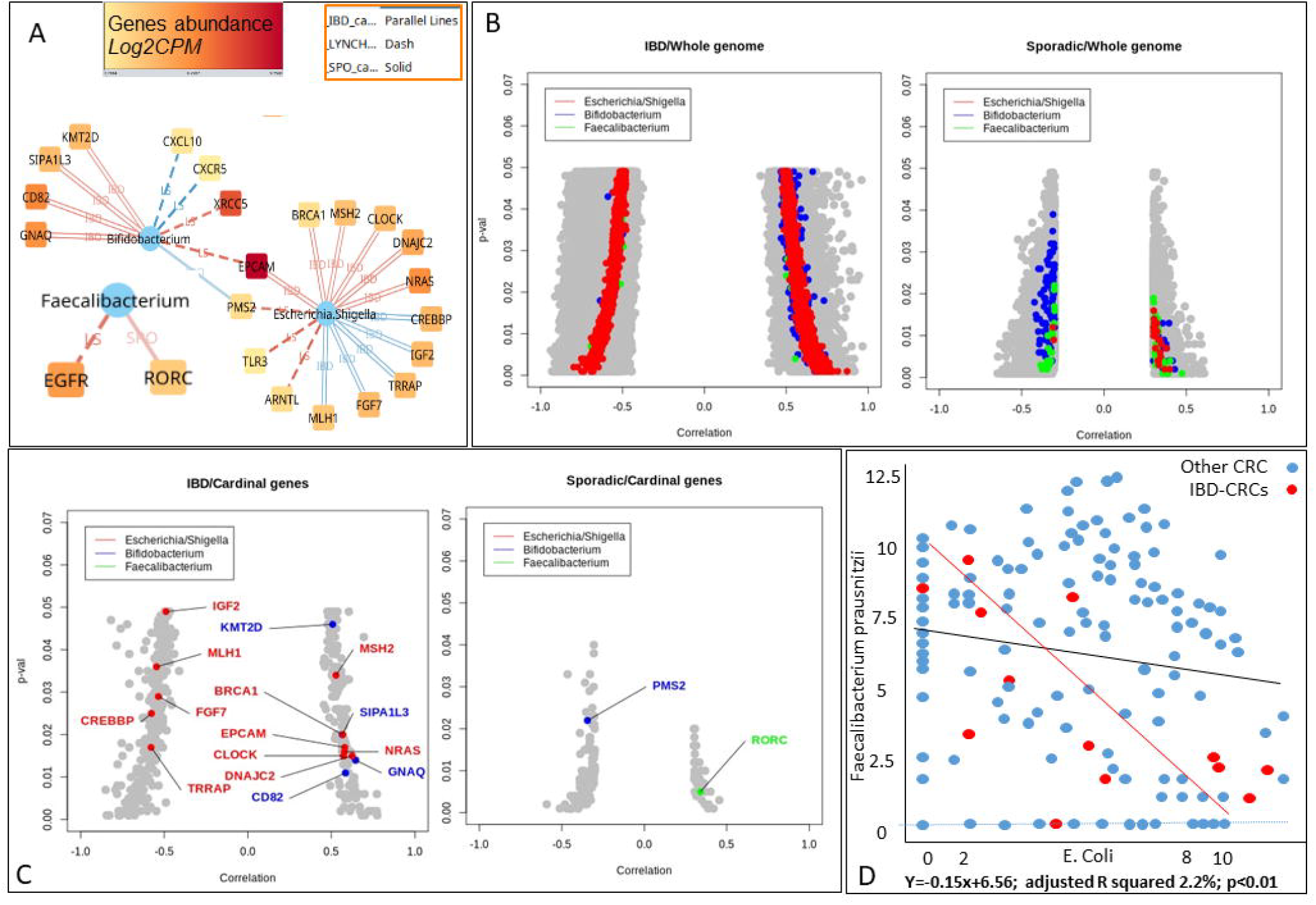
DNA (tumour, normal neighbor) and RNA from tumour tissues were extracted and submitted to 16sRNA and RNA sequencing (Illumina technology), respectively. Results were analyzed using Shaman platform and R program. Any correlation between two genes is shown using grey circle. Coloured (red, beu, green) circles design correlation between bacteria (red, Escherichia; bleu, Bifidobacterium; green, Faecalibacterium) and at least one gene with p-values set between 0.0001 and 0.05 and |correlation coefficients |>0.3; negative values design negative correlations and positive values positive correlations. (A) Correlation between PHLPP1 and 137 cardinal genes show different pattern according to CRC types with IBD displaying more negative correlations. (B) Taking all genes in consideration IBD (left) displays higher numbers of association than sporadic. (C) Focused on only 137 cardinal genes eleven genes were found associated to E. Coli in IBD (left) while no association was found in sporadic cases (right). (D) Escherichia Coli and Faecalibacterium Praunitzii species adherent to tissues show linear correlation in IBD-CRC patients (red) and different than in remaining patients (bleu).

Virulent/Symbiotic imbalance and associations with genes involved in inflammatory and immune pathways conducted to analyze PHLPP1 gene in inflammatory and immune cells by using RNAScope and Immunohistochemistry.

### CD8 and INF RNAscope screening in tumour tissues

RNAscope for gene expressions within tumour tissue as compared to normal neighbor showed larger heterogeneity in sporadic CRC patients than in IBD- or Lynch-CRCs. PHLPP1, was expressed in all cells within normal and tumour locations, with roughly lower expression in tumours than in normal neighboring tissues. PHLPP1 has been co-stained with CD8 and IFNγ(PHLPP1&CD8, and PHLPP1&IFNγ or triple stained PHLPP1&CD8&INFγ) and cells quantified (Figure 6). Numbers of CD8 cytotoxic and macrophages INFγ-producing cells with detectable PHLPP1 gene expression was significantly lower in IBD-CRC as compared to sporadic-CRC cases (Figure 6A-C).

**Figure 6.**
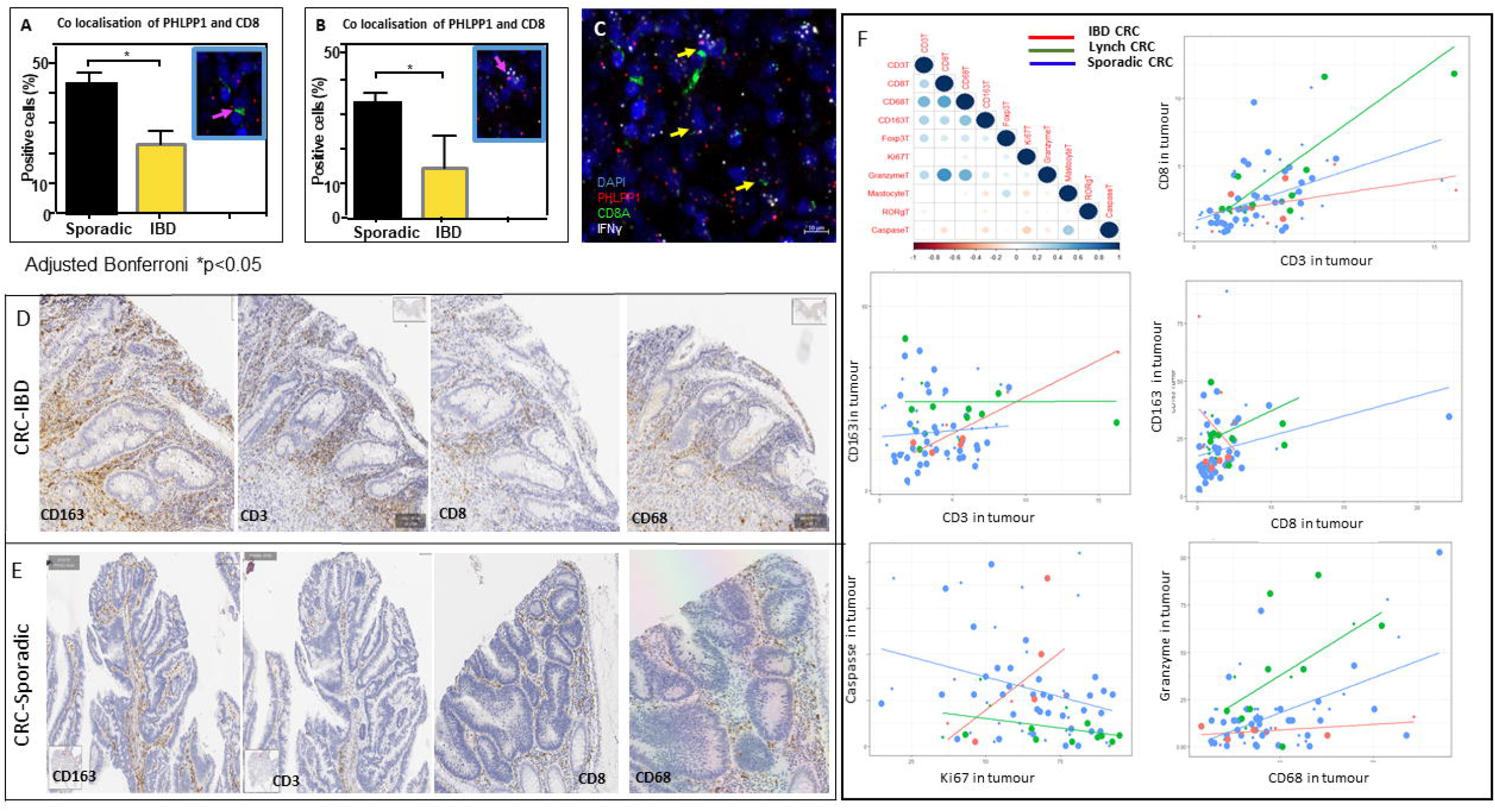
RNAscope In situ hybridization of inflammatory (IFNγ and CD8 (cytotoxic)) marker genes as well as PHLPP1 gene here taken as an anti-oncogene out of cardinal genes was performed on tumour tissue slides in sporadic and in IBD-CRCs. Note lower expression of PHLPP1 and CD8 as compared to sporadic cases in inflammatory and immune cells. A. high magnification of DAPI; B. Note lower expression of PHLPP1 spots in IBD-CRC tissues; C. multiples staining of three makers with significant differences for INF⍰ and CD8 co stained with PHLPP1. D and E show CD63, CD3, CD8 and CD68 immune stained cells. F shows correlations between various stained cells using Q Path morphometric program. Specific antibodies were purchased (Novogen; Biotechne, France) to identify Myeloid-macrophage CD68 and CD163 cells, Caspase 1 and Granzyme A as well as CD3, CD8 and quantified using QuPath program.

### Inflammatory and immune cell quantification in human tissues according to CRC types

To validate RNAScope results, we quantified inflammatory and immune cell infiltrates within tumor tissues using specific antibodies targeting myeloid and macrophage cells, CD68, CD163, CD3, CD4, CD8, FOXP3, ROR⍰T, and mastocytes in both normal and tumor serial tissue sections. The QuPath software program was employed for the morphometric quantification of immunostained cells (Figure 6D-F).

The normal colon tissues were infiltrated by CD3^+^ and CD4^+^, Granzyme-B^+^ and ROR⍰t^+^ FOXP3^+^ immune cells. The tumour tissues were distinguished from normal neighbor tissues by the diminution of Granzyme B^+^ CD3^+^ immune cells and ROR⍰t^+^FOXP3^+^ immune cells. Differences were dependent on types of CRCs (Lynch-, sporadic-, and IBD-associated). Notably, macrophages exhibited significantly higher infiltration in IBD-CRC tumors type (comparison between two significant cases: sporadic versus IBD). Furthermore, the patterns of co-associations such as “CD3-CD8”, and “CD3-macrophages (e.g. CD68, CD163)”, “CD8-macrophages”, and “CD8 with Granzyme-A or Caspase”, varied between CRC types with IBD-CRC tumours being significantly different than LS-CRC tumours (Figure 6F).

### Levels of PHLPP1 expression in IBD intestinal tissues

To validate our results regarding lower expression of PHLPP1 in tumour and inflamed no tumoral tissues versus normal tissues, two other series were investigated. The COAD cohort from the TCGA, for the levels of PHLPP1 RNA in CRC tumor versus normal tissues as well as in IBD patients’ colonic mucosa dataset to determine the influence of inflammatory injury on the expression of PHLPP1 (Figure S7). Despite broad variation of PHLPP1 gene expression in tumor tissues (n=475), there was a significant downregulation compared to neighboring normal tissues (n=41). In IBD patients’ intestinal tissues, levels of PHLPP1 lower at onset of the disease raised to normal values only in those patients responding to anti TNF-α therapy.

## DISCUSSION

This study examines the gene expression profiles, microbial adherence, and associated signaling pathways in CRC tissues, with a particular focus on IBD-associated CRC. We found that IBD-CRC is characterized by the downregulation of several tumor suppressor genes, particularly PHLPP1 gene that influences patients’ outcome. Along with IL17R, TRAPP, PHLPP1 gene plays a crucial role in the inflammatory response, spliceosome machinery, and autophagy. In agreement with this observation, gut analysis reveals that the imbalance between virulent and symbiotic bacteria is significantly more pronounced in IBD-CRC tissues compared to sporadic or Lynch syndrome-associated CRCs. This bacterial imbalance is closely associated with increased recruitment of inflammatory cells and the dysfunction of immune cells within the colonic tissue. The down expression of this gene is likely related to host epigenetic response to bacteria imbalance.

Based on global gene expression analysis, most of IBD-CRC cases were of CMS4 or CMS3 suggesting involvement of epithelial mesenchymal transition (EMT) activation^12,22,23^, where autophagy and inflammasome pathways are particularly activated. These pathways are strongly linked to innate immune system activation in response to various pathogens. In this regard, the cancer patient may develop various strategies that allow escape from these responses^24^. Adherent-invasive *E. Coli* (AIEC) has been reported to impede autophagy clearance in Crohns’ Diseasae (CD)^5,25^ but its’ mechanism of action is elusive.

The number of genes showing a significantly negative correlation with the PHLPP1 gene in IBD cases was three times higher than in sporadic CRC tissues, with the majority being involved in the spliceosome pathway. This pathway has been shown to be up regulated through epigenetic mechanisms in response to *Bacteroides fragilis*, a virulent bacteria which is strongly associated with the occurrence of IBD and in colitis-associated-CRC^26^. In the present series, we identified three bacterial genera (*Porphyromonas, Haemophilus, Pseudomonas)* and two species (*Porphyromonas perfringens* and *Shigella bonydii)* that were significantly overabundant in tumor tissues. These bacteria can be added to the earlier described CRC causing *Fusobacteria, Porphyromonas, or Parvimonas* genera^27,28^. Indeed, *P. micra* strains can directly alter the DNA methylation profile of promoters of key tumor-suppressor genes including those involved in EMT^29^.

Virulent bacteria whose adhesion to the tumour tissue impacts host homeostasis can alter the expression of genes involved in inflammatory and immune responses^30,31,32,33,34,35,36,37^. Down regulation of anti-oncogenes such as PHLPP1 gene in tumours of the present IBD-CRC cohort cannot be explained by exomic mutations (Table S5, suppl file); hence, we suggest they are epigenetically down-regulated in IBD patients as commonly reported for genes in inflamed tissues^38,39^. This is consistent with results from the IBD cohort dataset showing that expression of PHLPP1 recovers to the normal levels once the inflammation is controlled by anti TNF-α therapy (Figure S7, suppl file). Consistent with our findings, it has been reported that over a thousand differentially methylated CpG sites in the circulating DNA of IBD patients returned to normal levels and correlated with plasma C-reactive protein levels in those who responded to anti-inflammatory therapies^40^. Methylation of host genes prior to onset of tumor growth has been reported in IBD-cancer tissues, including SFRP1, SFRP2, SFRP5, TFPI2, Sox17, and GATA4 genes^9,39^. The downregulation of PHLPP1 in normal colonic epithelial cells contributes to the reduced activity of several pathways, including those involved in epithelial proliferation, motility, and inflammation^41,42^. PHLPP1 regulates both innate and adaptive immunity: in macrophages, it dampens signaling through TLR4 and the IFN-γ receptor inhibiting STAT1-mediated inflammatory genes and suppressing function of regulatory T cells^43^.

Dysregulated immune response due to gut dysbiosis might have been the cause of alteration in DNA methylation in T cells^24,44^. Thus, we speculate on the role of immunosuppressive medicine in IBD-CRC patients as a co-factor in explaining the IBD-CRC patients’ poor outcomes through immune cell failures. Immunosuppressive drugs can influence anti-tumour immune responses and restoration of PHLPP1 and negative regulation of AKT is expected after anti TNF-α therapy^45–47^.

T cells mediate anti-microbe and in the anti-cancer immune responses are the backbone of cancer immunotherapy^48^. In this regard, we observed a significant association between *Faecalibacterium* abundance, a well-known anti-inflammatory symbiotic, and RORC transcripts in sporadic CRC but not in IBD-CRC tumours. Regulatory T cell (Treg) activity through the T cell receptor and IL-2 receptor requires PHLPP1 mediated activation of the PI3K/Akt pathway^49^, while downregulation of this tumor suppressor genes is associated with pathways such as Infectious Diseases and Cancer Disease, Oxidative Phosphorylation, Thermogenesis, Mitophagy, and Proteolysis with Spliceosome being the major pathway affected in IBD-CRCs^20,21^. That PHLPP1, CD8, INF triple staining in tumour tissues showed immune cells with optimal PHLPP1 expression were significantly lower in IBD-CRC as compared to sporadic CRC tissues, suggest, virulent/symbiotic bacteria imbalance in IBD-CRC reflects a resilient regulatory process to prevent subverting the host immunity as reported in plants during over-abundance of pathogens^50^.

## CONCLUSION

We have shown that tumor adherent microbiota dysbiosis with imbalance between adherent virulent bacteria and commensals in CRC tumour tissues influences patients’ outcomes, in association with PHLPP1 expression likely due to an epigenetic pathway.

## Data Availability

The shotgun metagenomic sequencing data and the 16S rRNA amplicon sequencing data are available from the European Nucleotide Archive (ENA) database (http://www.ebi.ac.uk/ena) under the accession number ERP005534. Additional data related to this paper will be available from the European Nucleotide Archive (ENA) database (http://www.ebi.ac.uk/ena) under the accession nos. ERX3622297–ERX3622402, ERR3628499–ERR3628604, ERS3936180–ERS3936285, and PRJEB35144.

## Acknowledgement

URC APHP Hopital St Antoine, Hopital Henri Mondor; CRB Hopital Henri Mondor

## Clinical trial concerned

CCR1; ITEP (NCT00624260); VATNIMAD (NCT01270360);

## Ethical committee approval

under the number 09-2016 was obtained on March 14, 2016 and PHRC VATNIMAD (Clinical Trial NCT01270360). Written informed consent to participate was written in French and obtain prior to the enrollment and sampling.

## Consent for publication

Written informed consent for publication was obtained from everyone.

## Contribution of each author

Prove of concept and experimental design (IS, KK, DM); analysis of results (IS, DM, KK, AW, CC, NOA); Funding (IS, KK, MC); Material & Clinical contribution (EB, AA, IS, FB, CT, CC, MS); regulatory and control quality (IS, CB); Writing (IS, DM, KK) and editing (all authors).

