## Supplementary files for "Poor prognosis in IBD-complicated colon cancer through gut dysbiosis-related immune response failure"

### Figure S1

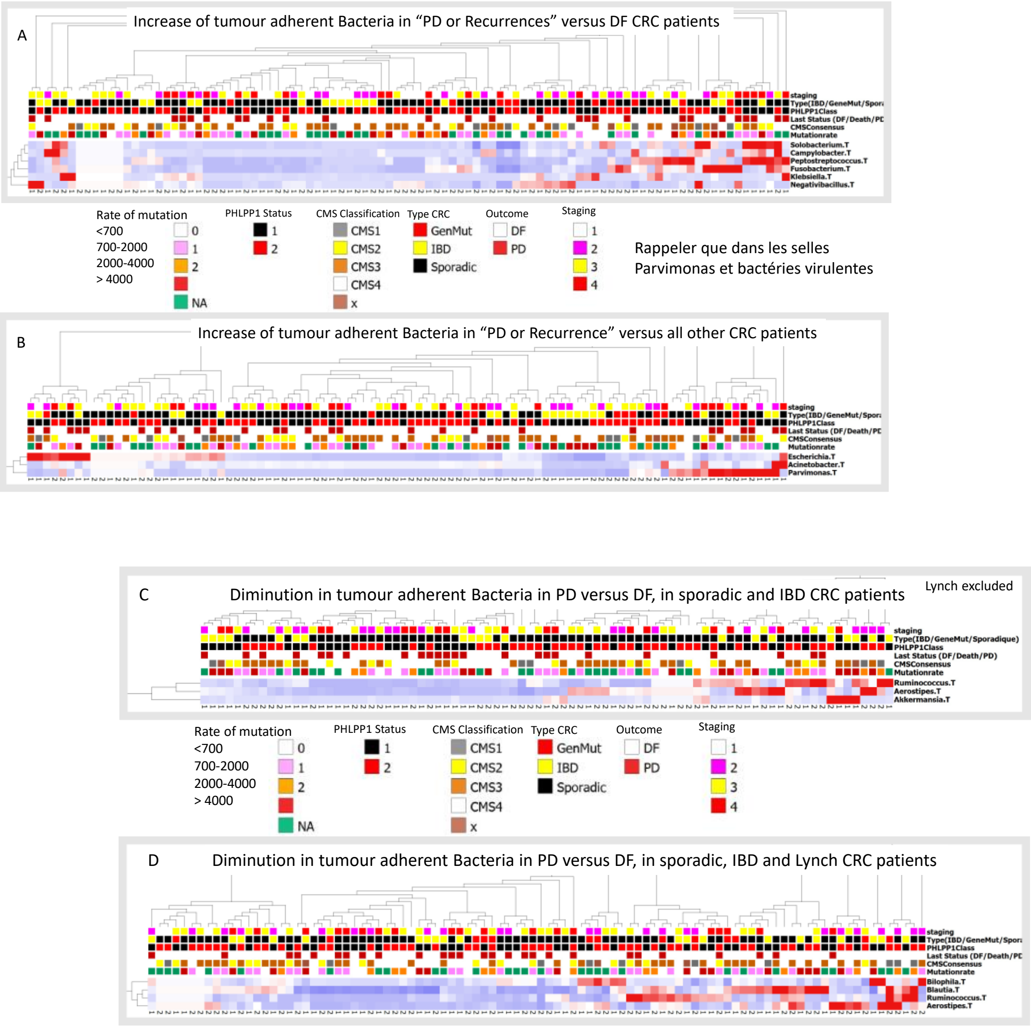

***Legends to figure S1***. *Adherent bacteria based on 16sRNA sequencing. which was performed after the prokaryote DNA extracts. Heatmap in a new analysis of bacteria in regard only to CRC patients displaying disease free survival (DFS) versus those with progression disease(PD)* ***in A*** *shows overabundance of several virulent bacteria when patients presenting with recurrence or progression of the disease were compared only to those with those remained free of disease.* ***B*** *shows same comparison when all patients (disease free plus stable disease) were considered Esherichia and Parvimonas overabundances might favor bad outcome.* ***C and D*** *show mainly diminution of symbionts in IBD. GenMut=Lynch syndrome; IBD=inflammatory bowel disease; CMS: Colon cancer Molecular Subtype; .T suffix after the name of bacteria design the microorganism adherent to tumoral tissue.*

**Figure S2.**

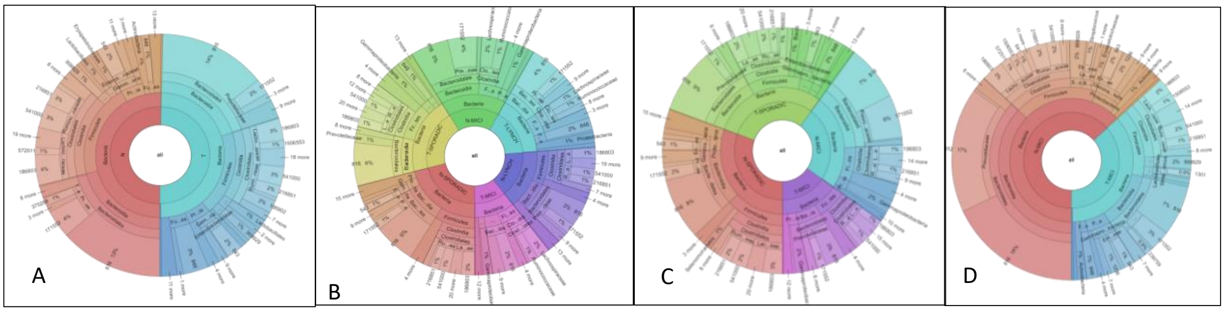

**Legends to figure S2.** Taking all samples and CRC types together. no significant difference was observed between N vs T regarding taxonomic features (A). although T cases appeared less diversified (B and C) whatehver the comparisons included all three types. Sporadic and IBD or IBD alone (D). Note that in IBD patients. loss of diversity in T is dramatic as compared to other types; T=tumour; N= normal neighbor; MICI=IBD

Korona pattern in three different CRC types [sporadic. IBD-MICI and Lynch syndrome] showed dramatic reduced diversity in tumour (T) in IBD (MICI) as compared to sporadic cases while there was similar pattern in tumour (T) and normal neighbor (N) when all patients were considered together.

**Table S1:**

**Species tissue adherent bacteria according to PHLPP1 and CRC types**

| | ID | Mean base | FoldChange | log2Fold | Adjusted p |
|---|---|---|---|---|---|
| Contrast PHLPP1 High vs Low | Klebsiella pneumoniae | 8.21 | 1.08e-01 | -3.205 | 0.036572 |
| | Shigella boydii | 7.99 | 1.98e-01 | -2.335 | 0.009600 |
| | Porphyromonas gingivalis | 70.97 | 1.01e-02 | -6.618 | 0.000000 |
| | Escherichia coli | 9.39 | 7.52e-02 | -3.732 | 0.000001 |
| Contrast Sporadic (Sp) vs IBD | Porphyromonas gingivalis | 70.97 | 2.49e+01 | 4.642 | 0.0458258 |
| | Criibacterium bergeronii | 151.78 | 6.38e+01 | 5.998 | 0.0190811 |
| | Bifidobacterium longum | 224.17 | 6.68e+00 | 2.742 | 0.0116255 |
| | Faecalicoccus pleomorphus | 6.24 | 8.58e+01 | 6.425 | 0.0116255 |
| | Shigella boydii | -7.99 | 2.1e+01 | 4.406 | 0.0003110 |
| | Lactococcus lactis | -8.61 | 4.96e+01 | 5.632 | 0.0000581 |
| IBD vs Lynch | Lactococcus lactis | -8.61 | 3.01e+01 | 4.912 | 0.049315 |
| Sp vs Lynch | Bifidobacterium longum | - 224.17 | 1.76e+01 | 4.139 | 0.000359 |
| | Streptococcus constellatus | 57.98 | 6.81e+01 | 6.092 | 0.001677 |
| | Faecalicoccus pleomorphus | 6.24 | 6.63e+01 | 6.052 | 0.044932 |

**Table S2: Data set analysis of tumour samples' RNA sequences for CMS assignation**[11.17]

| samplename | prediction. cmscaller | d.CMS1. cmscaller | d.CMS2. cmscaller | d.CMS3. cmscaller | d.CMS4. cmscaller | p.value. cmscaller | FDR. cmscaller |
|---|---|---|---|---|---|---|---|
| X0B03600201 | CMS4 | 0.72 | 0.68 | 0.70 | 0.60 | 0.00 | 0.00 |
| X0B03983701 | CMS4 | 0.67 | 0.76 | 0.65 | 0.57 | 0.00 | 0.00 |
| X0B12884101 | CMS3 | 0.73 | 0.73 | 0.69 | 0.78 | 0.00 | 0.00 |
| X0B13012701 | CMS4 | 0.63 | 0.76 | 0.71 | 0.58 | 0.00 | 0.00 |
| X1471051576 | CMS3 | 0.64 | 0.70 | 0.62 | 0.70 | 0.00 | 0.00 |
| X1800772624 | CMS3 | 0.66 | 0.75 | 0.55 | 0.66 | 0.00 | 0.00 |
| X1B07170901 | CMS3 | 0.72 | 0.80 | 0.68 | 0.77 | 0.00 | 0.00 |
| X2309190565 | CMS4 | 0.57 | 0.71 | 0.70 | 0.52 | 0.00 | 0.00 |
| X2380033088 | CMS2 | 0.75 | 0.66 | 0.79 | 0.74 | 0.01 | 0.01 |
| X2380034262 | CMS1 | 0.59 | 0.74 | 0.75 | 0.83 | 0.00 | 0.00 |
| X2510480101 | CMS2 | 0.76 | 0.60 | 0.70 | 0.62 | 0.00 | 0.00 |
| X2800243266 | CMS3 | 0.71 | 0.72 | 0.63 | 0.75 | 0.00 | 0.00 |
| X2800852568 | CMS1 | 0.72 | 0.77 | 0.74 | 0.79 | 0.03 | 0.03 |
| X2800993001 | CMS1 | 0.60 | 0.75 | 0.80 | 0.74 | 0.00 | 0.00 |
| X3309282085 | CMS3 | 0.61 | 0.73 | 0.60 | 0.80 | 0.00 | 0.00 |
| X3390029202 | CMS3 | 0.66 | 0.71 | 0.61 | 0.82 | 0.00 | 0.00 |
| X3400260101 | CMS4 | 0.74 | 0.80 | 0.81 | 0.52 | 0.00 | 0.00 |
| X360047160.T | CMS3 | 0.69 | 0.73 | 0.57 | 0.82 | 0.00 | 0.00 |
| X3800001753 | CMS4 | 0.64 | 0.74 | 0.65 | 0.55 | 0.00 | 0.00 |
| X4400033186 | CMS1 | 0.63 | 0.70 | 0.73 | 0.76 | 0.00 | 0.00 |
| X4400039331 | CMS2 | 0.68 | 0.64 | 0.67 | 0.85 | 0.01 | 0.01 |
| X4400048158 | CMS3 | 0.75 | 0.73 | 0.68 | 0.84 | 0.00 | 0.00 |
| X461023583 | CMS1 | 0.64 | 0.70 | 0.66 | 0.79 | 0.00 | 0.00 |
| X461024737 | CMS4 | 0.71 | 0.62 | 0.75 | 0.55 | 0.00 | 0.00 |
| X4800337146.T | CMS2 | 0.80 | 0.61 | 0.66 | 0.82 | 0.00 | 0.00 |
| X4800341673 | CMS2 | 0.78 | 0.62 | 0.71 | 0.85 | 0.00 | 0.00 |
| X4800349374 | NA | 0.68 | 0.67 | 0.72 | 0.75 | 0.36 | 0.37 |

| | | | | | | | |
|---|---|---|---|---|---|---|---|
| X4800374624 | CMS3 | 0.78 | 0.67 | 0.65 | 0.79 | 0.00 | 0.00 |
| X4800383870 | CMS2 | 0.76 | 0.59 | 0.72 | 0.81 | 0.00 | 0.00 |
| X4800393405 | CMS3 | 0.78 | 0.70 | 0.68 | 0.87 | 0.00 | 0.00 |
| X4800393826 | CMS4 | 0.58 | 0.76 | 0.71 | 0.58 | 0.00 | 0.00 |
| X4800415901 | NA | 0.71 | 0.69 | 0.74 | 0.78 | 0.98 | 1.00 |
| X4800422427 | CMS4 | 0.69 | 0.64 | 0.72 | 0.40 | 0.00 | 0.00 |
| X4800429512 | CMS4 | 0.69 | 0.70 | 0.80 | 0.46 | 0.00 | 0.00 |
| X4800437558 | CMS3 | 0.75 | 0.67 | 0.66 | 0.86 | 0.00 | 0.00 |
| X4800445736 | CMS1 | 0.61 | 0.74 | 0.75 | 0.81 | 0.00 | 0.00 |
| X4800453462 | CMS2 | 0.79 | 0.61 | 0.78 | 0.75 | 0.00 | 0.00 |
| X4800460683 | CMS3 | 0.63 | 0.74 | 0.54 | 0.74 | 0.00 | 0.00 |
| X4800491425 | CMS4 | 0.75 | 0.67 | 0.77 | 0.66 | 0.00 | 0.00 |
| X4800498256 | CMS1 | 0.55 | 0.77 | 0.69 | 0.75 | 0.00 | 0.00 |
| X4800526692 | CMS3 | 0.70 | 0.68 | 0.62 | 0.84 | 0.00 | 0.00 |
| X4800559056 | CMS4 | 0.66 | 0.79 | 0.67 | 0.52 | 0.00 | 0.00 |
| X4800576020 | CMS4 | 0.71 | 0.77 | 0.75 | 0.42 | 0.00 | 0.00 |
| X4800584221 | CMS1 | 0.60 | 0.81 | 0.76 | 0.80 | 0.00 | 0.00 |
| X4800594774 | CMS4 | 0.67 | 0.73 | 0.80 | 0.38 | 0.00 | 0.00 |
| X4800599680 | CMS4 | 0.73 | 0.59 | 0.75 | 0.52 | 0.00 | 0.00 |
| X4800621327 | NA | 0.67 | 0.66 | 0.73 | 0.74 | 0.05 | 0.05 |
| X4800628986 | CMS2 | 0.67 | 0.60 | 0.68 | 0.64 | 0.00 | 0.00 |
| X4800629587 | CMS2 | 0.79 | 0.64 | 0.78 | 0.71 | 0.00 | 0.00 |
| X4800638075 | CMS4 | 0.74 | 0.60 | 0.69 | 0.59 | 0.00 | 0.00 |
| X4800638653 | CMS4 | 0.70 | 0.80 | 0.84 | 0.56 | 0.00 | 0.00 |
| X4800674595 | NA | 0.79 | 0.71 | 0.70 | 0.90 | 1.00 | 1.00 |
| X4800674978 | CMS3 | 0.74 | 0.63 | 0.61 | 0.75 | 0.00 | 0.00 |
| X4800675124 | CMS4 | 0.71 | 0.69 | 0.76 | 0.44 | 0.00 | 0.00 |
| X4800685780 | CMS1 | 0.60 | 0.69 | 0.71 | 0.76 | 0.00 | 0.00 |
| X4800703082 | CMS3 | 0.66 | 0.67 | 0.63 | 0.84 | 0.00 | 0.00 |
| X4800706759 | CMS3 | 0.74 | 0.68 | 0.59 | 0.69 | 0.00 | 0.00 |
| X4800707768 | CMS2 | 0.76 | 0.58 | 0.66 | 0.79 | 0.00 | 0.00 |

| | | | | | | | |
|---|---|---|---|---|---|---|---|
| X504830101 | CMS4 | 0.65 | 0.79 | 0.70 | 0.46 | 0.00 | 0.00 |
| X5319020203 | CMS2 | 0.80 | 0.60 | 0.74 | 0.79 | 0.00 | 0.00 |
| X5410047094 | CMS3 | 0.71 | 0.68 | 0.62 | 0.88 | 0.00 | 0.00 |
| X5505750101 | CMS4 | 0.66 | 0.78 | 0.65 | 0.55 | 0.00 | 0.00 |
| X5800171786 | CMS1 | 0.63 | 0.78 | 0.75 | 0.74 | 0.00 | 0.00 |
| X5800175154 | CMS2 | 0.78 | 0.59 | 0.73 | 0.70 | 0.00 | 0.00 |
| X5800192593 | CMS4 | 0.70 | 0.62 | 0.78 | 0.55 | 0.00 | 0.00 |
| X5800251166 | CMS3 | 0.75 | 0.73 | 0.59 | 0.87 | 0.00 | 0.00 |
| X5800255523 | CMS3 | 0.73 | 0.70 | 0.58 | 0.76 | 0.00 | 0.00 |
| X5800285558 | CMS4 | 0.75 | 0.80 | 0.81 | 0.54 | 0.00 | 0.00 |
| X5800877134 | CMS2 | 0.77 | 0.60 | 0.78 | 0.68 | 0.00 | 0.00 |
| X6329052481 | CMS4 | 0.72 | 0.66 | 0.72 | 0.50 | 0.00 | 0.00 |
| X6420006503_49 | CMS4 | 0.66 | 0.80 | 0.76 | 0.44 | 0.00 | 0.00 |
| X6420008956 | CMS2 | 0.78 | 0.63 | 0.75 | 0.80 | 0.00 | 0.00 |
| X6420010983_30 | CMS3 | 0.61 | 0.67 | 0.61 | 0.75 | 0.00 | 0.00 |
| X6420025424 | CMS3 | 0.75 | 0.69 | 0.60 | 0.87 | 0.00 | 0.00 |
| X6420059784 | CMS2 | 0.79 | 0.54 | 0.77 | 0.64 | 0.00 | 0.00 |
| X6800256222 | CMS4 | 0.72 | 0.64 | 0.81 | 0.55 | 0.00 | 0.00 |
| X6800282206 | CMS4 | 0.68 | 0.74 | 0.70 | 0.60 | 0.00 | 0.00 |
| X6800299931 | CMS3 | 0.71 | 0.68 | 0.62 | 0.77 | 0.00 | 0.00 |
| X6800808321 | NA | 0.67 | 0.77 | 0.69 | 0.87 | 0.07 | 0.07 |
| X7339027551 | CMS3 | 0.80 | 0.65 | 0.60 | 0.86 | 0.00 | 0.00 |
| X7339047662 | CMS3 | 0.76 | 0.67 | 0.62 | 0.76 | 0.00 | 0.00 |
| X7430012015 | NA | 0.73 | 0.70 | 0.71 | 0.71 | 0.07 | 0.08 |
| X7430029683_52 | CMS1 | 0.58 | 0.80 | 0.68 | 0.72 | 0.00 | 0.00 |
| X7430033675 | CMS4 | 0.69 | 0.73 | 0.80 | 0.53 | 0.00 | 0.00 |
| X7430053376 | CMS4 | 0.65 | 0.77 | 0.69 | 0.53 | 0.00 | 0.00 |
| X7517630101 | CMS3 | 0.66 | 0.76 | 0.62 | 0.69 | 0.00 | 0.00 |
| X8308129581 | CMS2 | 0.68 | 0.64 | 0.68 | 0.80 | 0.00 | 0.00 |
| X8308145823 | NA | 0.74 | 0.72 | 0.73 | 0.81 | 0.16 | 0.17 |
| X8308146995 | CMS4 | 0.73 | 0.68 | 0.65 | 0.57 | 0.00 | 0.00 |

| | | | | | | | |
|---|---|---|---|---|---|---|---|
| X8308174416 | CMS4 | 0.72 | 0.72 | 0.70 | 0.68 | 0.00 | 0.00 |
| X8440019869 | CMS4 | 0.60 | 0.74 | 0.69 | 0.44 | 0.00 | 0.00 |
| X8542010101 | NA | 0.73 | 0.71 | 0.72 | 0.74 | 1.00 | 1.00 |
| X8800964015_40 | CMS2 | 0.80 | 0.58 | 0.75 | 0.78 | 0.00 | 0.00 |
| X9308265017 | CMS3 | 0.75 | 0.68 | 0.57 | 0.73 | 0.00 | 0.00 |
| X9359049778 | CMS4 | 0.76 | 0.59 | 0.75 | 0.57 | 0.00 | 0.00 |
| X9543110101 | CMS1 | 0.58 | 0.78 | 0.62 | 0.75 | 0.00 | 0.00 |

**List S3: Cardinal genes involved in Colon Carcinogenesis**

"AKT1". "APC". "ARID1A". "ARNTL". "ARNTL2". "ATG16L1". "ATM". "ATR". "AXIN1". "BAX". "BRAF". "BRCA2". "CACNA1G". "CARD11". "CCND1". "CD82". "CDH1". "CLOCK". "COL16A1". "COX2". "CREBBP". "CTNNB1". "CX3CL1". "CXCL10". "CXCL9". "CXCR5". "DNAJC2". "E2F4". "EGFR". "EPCAM". "EPHA3". "ERBB2". "FBXW7". "FOXL2". "FOXP3". "GATA3". "GNA11". "GNAQ". "GZMA". "GZMB". "IGF2". "IL10". "IL17A". "IL17RA". "IL17RB". "IL17RD". "IL23A". "IL23R". "KIT". "KMT2D". "KRAS". "LAMB2". "MAP3K3". "MCPH1". "MED12L". "MET". "MGMT". "MLH1". "MMP26". "MSH2". "MSH6". "MYC". "MYCBP2". "NEUROG1". "NF1". "NLRP6". "NOD2". "NRAS". "PDGFRA". "PHLPP1". "PIK3CA". "PMS2". "PMS2P4". "PRF1". "PTEN". "PTGS2". "PTPN11". "RASGRP3". "RB1". "RET". "ROBO1". "RORC". "RUNX3". "PTPN11". "SIPA1L3". "SMAD4". "SOCS1". "TBX21". "TCF4". "TGFBR1". "TGFBR2". "TLR1". "TLR2". "TLR3". "TLR4". "TLR5". "TLR6". "TLR7". "TLR8". "TLR9". "TOPORS". "TP53". "TRAPPC6B". "TRRAP". "VEGFA". "FGFR1OP". "FGFBP3". "FGF1". "FGF2". "FGF3". "FGF4". "FGF5". "FGF6". "FGF7". "FGF8". "FGF9". "FGF10". "FGF11". "FGF12"."FGF13". "FGF14". "FGFR1". "FGFR3". "FGFR2". "FGFR4". "FGFR1OP2". "FGF20". "FGF21". "FGF22". "FGFRL1". "FGF23". "FGFBP2". "FGF18". "FGF17". "FGF16". "FGF19". "FGFBP1"

*The genes have been identified to be cardinal in colon carcinogenesis through scientific and experimental data in the literature*[23.53]

**Figure S3**

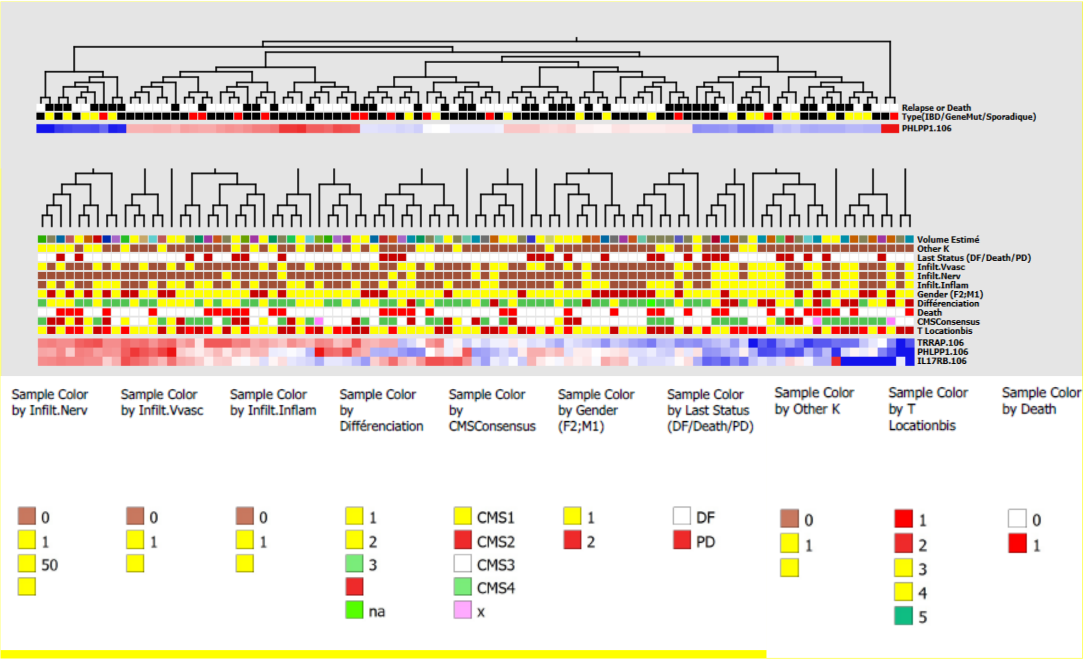

**Legends to figure S3: Heatmap from cardinal gene RNA levels in tumour tissues at baseline influencing patients' outcome (success. Failure)** *T*umour samples from n=96 CRC patients (n=56 sporadic; n=11 IBD and n=9 Lynch syndrome) were submitted to RNA extract and gene expression were used for clustering genes through an unsupervised analysis based on 137 cardinal genes; *t-test [<] was used comparing gene RNA values according to Recurrence/Progression diseases (DF) or disease free (DF) survival (1 vs 0; p = 0.002. q = 0.186; Eliminated factors None Filter by Standard Deviation (s/smax) ; Normalization Mean=0. Var=1 Missing value=1 ; Reconstruction N/A Collapse mode none Logarithm; two patients could not be analyzed due to material or technical issues. The heatmap shows levels of PHLPP1 gene RNA regarding CRC types and DF versus Recurrence/Death or the progression of the disease-PD. The Qlucore Omics Explorer uses structure data from an experiment to verify that the expected subgroups of subjects could be identified with variance filtering being used to reduce the noise. and the projection score to set the filtering threshold. The "Mean=0. Var=1" setting has been used to scale the data. No variable was left after the filtering. After filtering out variables with low overall variance to reduce the impact of noise. and centering and scaling the remaining variables to zero mean and unit variance. the projection score*[19] *was used to determine the optimal filtering threshold. retaining all variables. The identification of significantly differential variables between Low and high gene RNA subgroups was performed by fitting a linear model for each variable with gene level class as a predictor and including the sample factor as a nuisance covariate. P-values were adjusted for multiple testing using the Benjamini-Hochberg method*[15]*. and variables with adjusted p-values below 0.1 were considered significant. This resulted in significant variables.*

**Table S4. Cardinal Genes correlated with PHLPP1 gene according to CRC type.**

| | ne | | IBD | | LYNCH | | SPORADIC | | |
|---|---|---|---|---|---|---|---|---|---|
| Gene name | meanLog2CP | sd | corr | P val | corr | P val | corr | P val | CorrSTATUS |
| AXIN1 | 5.96 | 0.87 | 0.51 | 0.03 | 0.83 | 0.00 | 0.52 | 0.00 | All Cohorts |
| CREBBP | 5.48 | 1.07 | 0.74 | 0.00 | 0.82 | 0.00 | 0.58 | 0.00 | All Cohorts |
| SIPA1L3 | 5.14 | 1.05 | 0.58 | 0.01 | 0.82 | 0.01 | 0.59 | 0.00 | All Cohorts |
| TRRAP | 5.52 | 1.05 | 0.51 | 0.03 | 0.74 | 0.03 | 0.66 | 0.00 | All Cohorts |
| ARID1A | 5.59 | 1.04 | 0.48 | 0.04 | 0.72 | 0.04 | 0.62 | 0.00 | All Cohorts |
| KMT2D | 5.88 | 1.50 | 0.50 | 0.03 | 0.76 | 0.02 | 0.67 | 0.00 | All Cohorts |
| ATG16L1 | 3.04 | 1.77 | 0.03 | 0.91 | 0.75 | 0.03 | 0.60 | 0.00 | LYNCH-SPO |
| MGMT | 4.70 | 1.25 | -0.19 | 0.45 | -0.74 | 0.04 | -0.43 | 0.00 | LYNCH-SPO |
| CTNNB1 | 8.23 | 0.72 | -0.29 | 0.24 | 0.80 | 0.02 | 0.36 | 0.00 | LYNCH-SPO |
| ERBB2 | 7.33 | 0.93 | 0.37 | 0.13 | 0.70 | 0.04 | 0.48 | 0.00 | LYNCH-SPO |
| KRAS | 5.39 | 0.72 | -0.32 | 0.21 | 0.76 | 0.02 | 0.47 | 0.00 | LYNCH-SPO |
| SMAD4 | 5.54 | 0.73 | 0.38 | 0.12 | 0.85 | 0.01 | 0.30 | 0.01 | LYNCH-SPO |
| TP53 | 5.79 | 1.07 | -0.05 | 0.83 | 0.88 | 0.00 | 0.43 | 0.00 | LYNCH-SPO |
| BRCA1 | 3.64 | 1.68 | -0.66 | 0.00 | 0.48 | 0.23 | 0.46 | 0.00 | IBD-SPO |
| CHEK2 | 4.00 | 1.05 | -0.64 | 0.00 | -0.28 | 0.51 | 0.44 | 0.00 | IBD-SPO |
| LAMB2 | 6.49 | 1.48 | 0.71 | 0.00 | 0.01 | 0.98 | 0.32 | 0.01 | IBD-SPO |
| MSH6 | 4.96 | 0.76 | -0.50 | 0.03 | 0.18 | 0.68 | 0.38 | 0.00 | IBD-SPO |
| MAP3K3 | 4.25 | 1.05 | 0.63 | 0.01 | 0.81 | 0.01 | 0.26 | 0.03 | IBD-LYNCH |
| AKT1 | 7.14 | 0.84 | 0.21 | 0.40 | 0.62 | 0.10 | 0.59 | 0.00 | SPO |
| E2F4 | 6.15 | 0.78 | 0.01 | 0.97 | 0.57 | 0.14 | 0.58 | 0.00 | SPO |
| EGFR | 5.13 | 0.95 | 0.41 | 0.09 | 0.66 | 0.08 | 0.54 | 0.00 | SPO |
| FBXW7 | 4.40 | 1.04 | -0.07 | 0.79 | 0.68 | 0.06 | 0.58 | 0.00 | SPO |
| NF1 | 5.50 | 0.74 | 0.16 | 0.53 | 0.56 | 0.15 | 0.53 | 0.00 | SPO |
| PTPN11 | 6.14 | 0.93 | -0.03 | 0.92 | 0.58 | 0.13 | 0.53 | 0.00 | SPO |
| VEGFA | 7.70 | 1.59 | 0.06 | 0.80 | 0.55 | 0.16 | 0.55 | 0.00 | SPO |
| ATR | 4.70 | 0.81 | -0.31 | 0.21 | -0.39 | 0.34 | 0.53 | 0.00 | SPO |
| MYCBP2 | 4.69 | 1.21 | 0.06 | 0.82 | 0.23 | 0.58 | 0.53 | 0.00 | SPO |
| APC | 3.93 | 0.89 | 0.29 | 0.24 | 0.68 | 0.07 | 0.36 | 0.00 | SPO |
| GNA11 | 6.62 | 0.84 | 0.44 | 0.07 | 0.63 | 0.09 | 0.42 | 0.00 | SPO |
| IL17RA | 4.66 | 0.88 | 0.46 | 0.06 | 0.52 | 0.19 | 0.44 | 0.00 | SPO |
| NOD2 | 1.32 | 1.82 | 0.21 | 0.40 | 0.63 | 0.10 | 0.30 | 0.01 | SPO |
| TOPORS | 4.03 | 0.82 | -0.23 | 0.37 | 0.70 | 0.05 | 0.34 | 0.00 | SPO |
| ARNTL | 2.68 | 1.28 | 0.12 | 0.64 | 0.12 | 0.79 | 0.42 | 0.00 | SPO |
| ATM | 5.14 | 1.08 | 0.10 | 0.70 | -0.16 | 0.70 | 0.49 | 0.00 | SPO |
| BRAF | 4.55 | 0.67 | 0.39 | 0.11 | 0.47 | 0.25 | 0.50 | 0.00 | SPO |
| BRCA2 | 2.97 | 1.56 | -0.47 | 0.05 | 0.23 | 0.59 | 0.46 | 0.00 | SPO |
| CD82 | 5.81 | 0.93 | -0.12 | 0.65 | -0.17 | 0.70 | 0.33 | 0.01 | SPO |
| FGFR3 | 4.74 | 1.75 | 0.06 | 0.81 | 0.09 | 0.84 | 0.31 | 0.01 | SPO |
| MCPH1 | 3.04 | 1.16 | -0.26 | 0.30 | 0.29 | 0.49 | 0.34 | 0.01 | SPO |
| MYC | 6.34 | 1.49 | -0.13 | 0.59 | 0.27 | 0.53 | 0.38 | 0.00 | SPO |
| PMS2 | 3.03 | 1.10 | -0.16 | 0.53 | -0.03 | 0.94 | 0.41 | 0.00 | SPO |
| SOCS1 | 2.92 | 1.47 | 0.15 | 0.55 | 0.29 | 0.48 | 0.32 | 0.01 | SPO |
| CDH1 | 7.78 | 1.90 | -0.19 | 0.46 | 0.83 | 0.01 | 0.30 | 0.01 | LYNCH |
| GNAQ | 5.98 | 0.61 | 0.32 | 0.20 | 0.94 | 0.00 | 0.21 | 0.09 | LYNCH |
| PTEN | 4.46 | 0.86 | -0.34 | 0.17 | 0.83 | 0.01 | 0.17 | 0.16 | LYNCH |
| RORC | 3.85 | 1.76 | 0.14 | 0.57 | 0.77 | 0.01 | 0.11 | 0.37 | LYNCH |

| | | | | | | | | | |
|---|---|---|---|---|---|---|---|---|---|
| XRCC6 | 7.66 | 0.60 | -0.30 | 0.22 | 0.77 | 0.03 | 0.10 | 0.41 | LYNCH |
| CHEK1 | 4.08 | 0.98 | -0.70 | 0.00 | 0.52 | 0.19 | 0.24 | 0.04 | IBD |
| COL16A1 | 6.09 | 1.63 | 0.62 | 0.01 | -0.54 | 0.17 | 0.13 | 0.27 | IBD |
| FGFR1 | 4.24 | 1.89 | 0.76 | 0.00 | 0.59 | 0.13 | 0.24 | 0.05 | IBD |
| IGF2 | 3.63 | 3.04 | 0.63 | 0.01 | 0.58 | 0.13 | 0.07 | 0.56 | IBD |
| IL17RD | 2.35 | 1.83 | 0.50 | 0.03 | 0.50 | 0.22 | 0.28 | 0.02 | IBD |
| MSH2 | 4.35 | 1.06 | -0.63 | 0.01 | 0.36 | 0.37 | 0.26 | 0.03 | IBD |
| CX3CL1 | 3.56 | 1.40 | 0.52 | 0.02 | 0.25 | 0.55 | 0.14 | 0.25 | IBD |
| DNAJC2 | 5.49 | 0.78 | -0.54 | 0.02 | -0.07 | 0.86 | 0.20 | 0.10 | IBD |
| EPCAM | 9.49 | 2.02 | -0.46 | 0.03 | 0.49 | 0.22 | 0.10 | 0.40 | IBD |
| FGFR2 | 3.30 | 1.78 | 0.60 | 0.01 | 0.20 | 0.63 | 0.19 | 0.13 | IBD |
| NRAS | 5.83 | 0.72 | -0.63 | 0.01 | 0.04 | 0.93 | 0.08 | 0.50 | IBD |
| XRCC5 | 7.47 | 0.70 | -0.59 | 0.01 | 0.29 | 0.50 | 0.21 | 0.08 | IBD |

*RNAs have been extracted from tumour tissue samples and submitted to RNAseq; genes' RNA levels were compared. IBD:inflammatory bowel disease; sd: standard deviation; corr: correlation coefficient; CorrSTATUS: correlation was significant in SPI: sporadic. LYNCH: Lynch syndrome or ALL subtypes of CRCS; these genes have been identified as significantly correlated with PHLPP1 gene through a panel of cardinal genes as identified in the litterature[23].[53]*

**Figure S4**

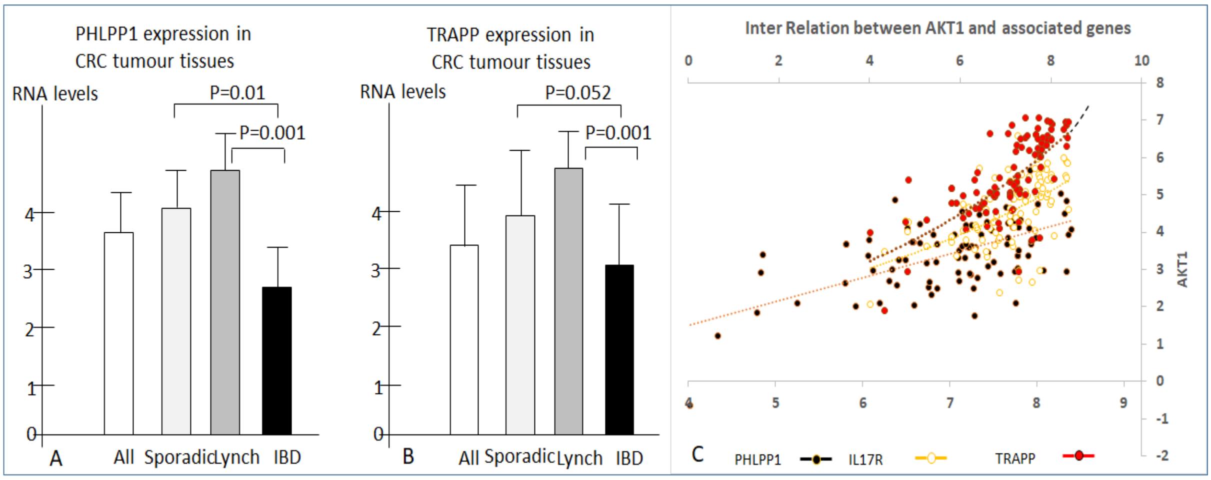

***Legends to figure S4****. RNA was extracted from tumour tissues and levels of gene. TRAPP and IL17R were found influencing patients' outcome (failure versus success) over a 3-yr period follow up. PHLPP1 (A) and TRAPP (B) were lower in IBD-CRC tumours tissues; when these genes were correlated with AKT1. PHLPP1 gene was the main pivotal with an exponential relationship (C).*

### Figure S5

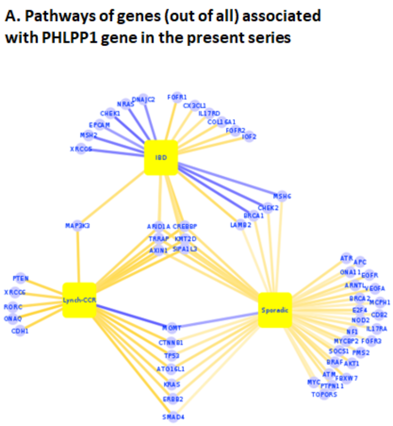

**B. Pathways of genes (out of all) associated with PHLPP1 gene in the present series**

| Signaling Pathway | Enrichment |
|---|---|
| Infection Disease | 20 |
| Cancer Disease | 17 |
| mTOR/TNF/TH1/2/17/PDL1/Rap1 | 13 |
| Degradation Protein/Sugar/Lipids | 12 |
| Degenerative /Senescence | 9 |
| Lipids/Phospholipids/Phosphorylation | 8 |
| Platelet/Vascular/Adhesin/Cytoskeleton | 6 |
| Muscle Actions/Apelin/ | 6 |
| Metabolism/Energy/Insulin | 5 |
| Sugar metabolism | 3 |
| Autophagy/Mitophagy | 2 |

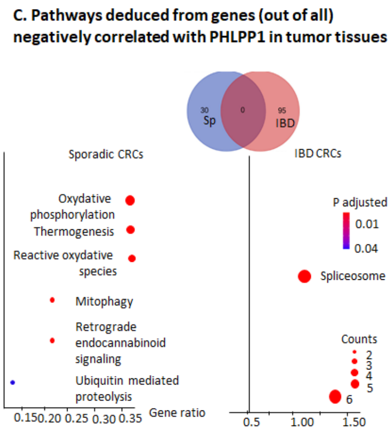

***Legends to figure S5***: **Genes correlations with PHLPP1 gene and networks**. *Several genes' RNA levels were found significantly correlated with PHLPP1 RNA. RNA was extracted from tumour tissue samples using Qiagen kit and materials were submitted to the sequencing by Illumina Hiseq technic.* ***A:*** *Based on RNA levels. HPLPP1 gene is differently co-related with other cardinal genes according to various CRC types.* ***B:*** *Genes positively correlated with PHLPP1 gene out of all are analysed by Enrich R and hierarchy of various pathways is indicated with number of genes mentioned to be enriched in pathways.* ***C.*** *Three times more genes were negatively correlated with PHLPP1 gene in IBD-CRC than in Sporadic-CRC cases and no one in Lynch syndrome.*

### Figure S6

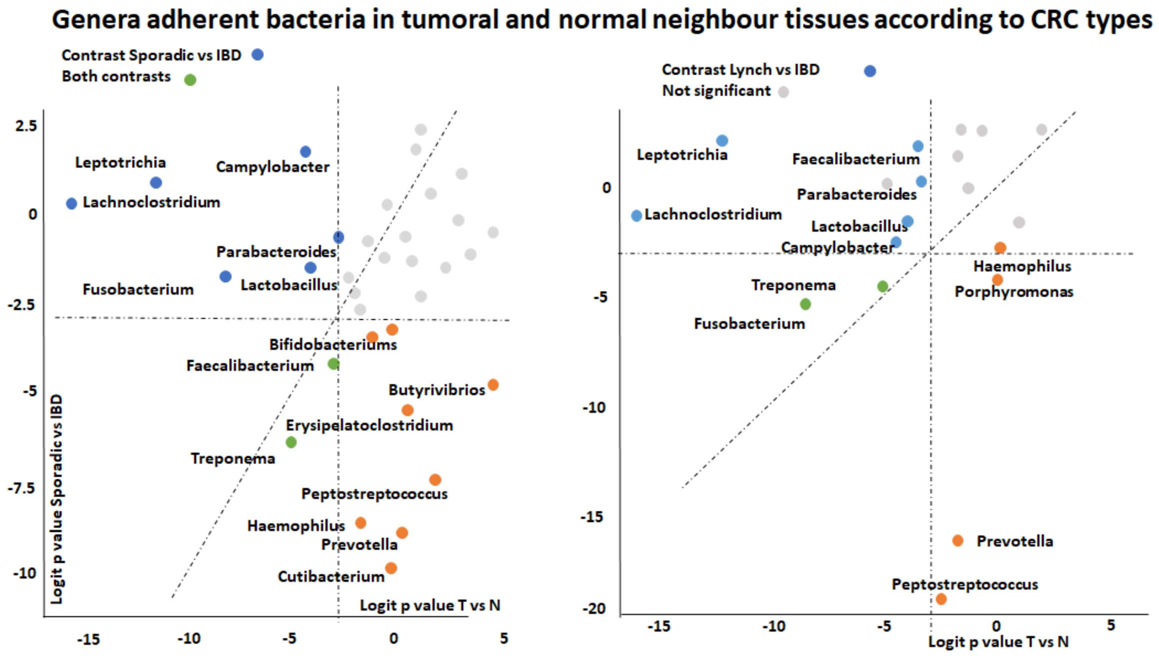

***Legends to figure S6***. **Differential bacteria adherent to tissues.** *DNA from (tumour. normal neighbor) tissues were extracted and submitted to 16sRNA sequencing procedure (Illumina technology). Results were analyzed using Shaman platform. Contrasts regarding genera adherent to tissues and CRC types (IBD vs Ly or Sporadic) revealed differential features.*

**Figure S7**

**A.PHLPP1 expressionin Normal and in Tumor tissues (TCGA dataset)**

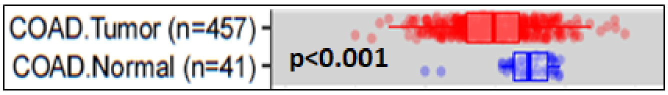

**B. PHLPP1 RNA Levels in IBD patients' intestinal tissues (IBD dataset)**

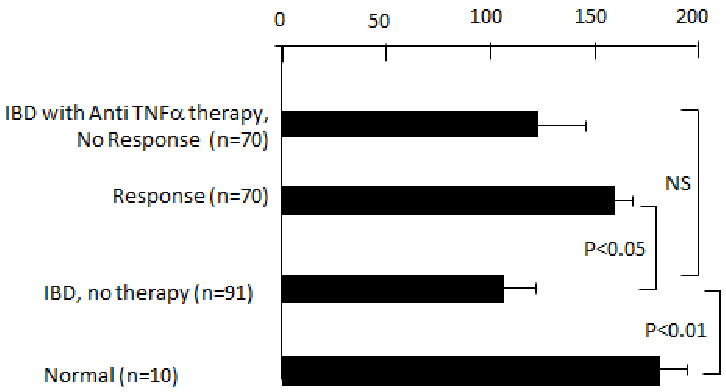

**Legends to figure S7. PHLPP1 expression in tumour and inflamed tissues**. Levels of PHLPP1 RNA in the colonic mucosa have been loaded from online available COAD dataset (*https://omicsview.org/human/app/gene_expressions/app_dashboard*) including adenocarcinoma (n=457) and normal colonic tissues (n=41) were compared and showed significantly lower RNA levels in tumour than in normal tissues (p<0.01) although large diversity and heterogeneity in cancer tissues were observed (A). Levels of PHLPP1 RNA in IBD patients' colonic tissue samples were obtained from IBD data set (*for raw data connect* ulcerative colitis and Crohn diseases through trial ID GSE23597. GSE12251. GSE52746. GSE16879) and showed PHLPP1 gene in patients with acute onset of disease was downregulated as compared to normal tissues and return to normal values only in patients responsive to anti TNF therapy (B).

**Table S5. Gene mutation/variants in genes clustering with PHLPP1 gene in colon or rectal tumour tissues.**

| | | | | | | |
|---|---|---|---|---|---|---|
| X2800243266 | PHLPP1 | missense_variant | | c.4507C> | p.Leu1503Val | SPORADIC |
| X2800243266 | | | | c.2971C> | p.Leu991Val | |
| X2800852568 | PHLPP1 | synonymous_variant | | c.5145G> | p.Thr1715Thr | SPORADIC |
| | | | | c.3609G> | p.Thr1203Thr | |
| X3309282085 | PHLPP1 | missense_variant | | c.4883C> | p.Ala1628Val | SPORADIC |
| | | | | c.3347C> | p.Ala1116Val | |
| X4400039331 | PHLPP1 | synonymous_variant | | c.2121A> | p.Pro707Pro | SPORADIC |
| | | | | c.585A>T | p.Pro195Pro | |
| X4400039331 | PHLPP1 | non_coding_transcript_exor | | n.642A>T | NA | SPORADIC |
| X4800341673 | PHLPP1 | intron_variant | | c.3324+32 | NA | SPORADIC |
| | | | | c.1788+32T>G | | |
| | | | | c.492+32T>G | | |
| | | | | n.*3393T>G | | |
| X4800415901 | PHLPP1 | intron_variant | | c.1112-20 | NA | LYNCH |
| | | intron_variant | | c.17-20_17-19delTT | | |
| | | intron_variant | | c.2648-20_2648-19delTT | | |
| | | 5_prime_UTR_variant | | c.-21dupG | | |
| | | 5_prime_UTR_variant | | c.-1557dupG | | |
| | | upstream_gene_variant | | n.-1083_-1082delTT | | |
| X4800548582 | PHLPP1 | synonymous_variant | | c.585A>T | p.Pro195Pro | SPORADIC |
| | | | | n.642A>T | NA | |
| | | | | c.2121A> | p.Pro707Pro | |
| X5800275405 | PHLPP1 | downstream_gene_variant | | n.*3189de | NA | LYNCH |
| | | | | c.1626-10delT | | |
| | | | | c.3162-10delT | | |
| | | | | c.330-10delT | | |
| X6420008956 | PHLPP1 | upstream_gene_variant | | n.-1083de | NA | SPORADIC |
| | | | | c.2648-20delT | | |
| | | | | c.1112-20delT | | |
| | | | | c.17-20delT | | |
| X6800299931 | PHLPP1 | intron_variant | | n.155-22_ | NA | SPORADIC |
| | | | | n.1085_1086delTC | | |
| | | | | n.368-22_368-21delTC | | |
| | | | | n.169-22_169-21delTC | | |
| | | | | c.1774-22_1774-21delTC | | |
| | | | | c.238-22_238-21delTC | | |
